# Dissociated development of speech and limb sensorimotor learning in stuttering: speech auditory-motor learning is impaired in both children and adults who stutter

**DOI:** 10.1101/2020.09.23.310797

**Authors:** Kwang S. Kim, Ayoub Daliri, J. Randall Flanagan, Ludo Max

## Abstract

Stuttering is a neurodevelopmental disorder of speech fluency. Various experimental paradigms have demonstrated that affected individuals show limitations in sensorimotor control and learning. However, controversy exists regarding two core aspects of this perspective. First, it has been claimed that sensorimotor learning limitations are detectable only in adults who stutter (after years of coping with the disorder) but not during childhood close to the onset of stuttering. Second, it remains unclear whether stuttering individuals’ sensorimotor learning limitations affect only speech movements or also unrelated effector systems involved in nonspeech movements. We report data from separate experiments investigating speech auditory-motor learning (*N* = 60) and limb visuomotor learning (*N* = 84) in both children and adults who stutter versus matched nonstuttering individuals. Both children and adults who stutter showed statistically significant limitations in speech auditory-motor adaptation with formant-shifted feedback. This limitation was more profound in children than in adults and in younger children versus older children. Between-group differences in the adaptation of reach movements performed with rotated visual feedback were subtle but statistically significant for adults. In children, even the nonstuttering groups showed limited visuomotor adaptation just like their stuttering peers. We conclude that sensorimotor learning is impaired in individuals who stutter, and that the ability for speech auditory-motor learning—which was already adult-like in 3-6 year-old typically developing children—is severely compromised in young children near the onset of stuttering. Thus, motor learning limitations may play an important role in the fundamental mechanisms contributing to the onset of this speech disorder.

## Introduction

Reaching a thorough understanding of the biological foundations of stuttering requires parallel investigations of the “distal” cause(s) (What causes an individual to have the disorder?) as well as the “proximal” cause(s) (What are the mechanisms that cause instances of stuttering moments in the individual’s speech?) (Packman & Attanasio, 2004). Several research groups’ recent accomplishments in the areas of neuroimaging (e.g., Beal et al., 2013; Cai et al., 2014; Chang et al., 2011; Choo et al., 2012; Sitek et al., 2016; Wymbs et al., 2013) and gene discovery (e.g., Benito-Aragón et al., 2020; Frigerio-Domingues & Drayna, 2017; Frigerio-Domingues et al., 2019; Kang et al., 2010; Raza et al., 2012) indicate substantial progress regarding the former of these two causality questions. Progress in addressing the latter question continues to lag as very few specific hypotheses have been formulated to suggest biologically plausible mechanisms that may explain the individual occurrences of sound/syllable repetitions and sound prolongations.

In previous theoretical work, we have suggested one potential proximal cause that may occur within the sensorimotor system itself as the direct result of a dysfunction in well-documented sensorimotor mechanisms (Max, 2004; Max et al., 2004; Neilson & Neilson, 1987). In particular, the proposed framework was based on psychophysical data and computational modeling suggesting that for voluntary movements the central nervous system (CNS) relies on both (a) a controller that determines which motor commands will achieve a desired movement outcome (sometimes referred to as a control policy, feedback control policy, or internal inverse model), and (b) a feedback system that uses efference copy and an internal forward model in combination with afferent information and prior knowledge to best estimate the system state (e.g., Desmurget & Grafton, 2000; Kawato, 1999; Krakauer & Mazzoni, 2011; Shadmehr et al., 2010; Wolpert & Miall, 1996; Wolpert & Flanagan, 2009). Internal models are stored neural representations of effector- and environment-dependent mappings between motor commands and their sensory consequences. An individual’s ability for sensorimotor learning can be tested with adaptation paradigms in which the system’s input-output mapping is experimentally manipulated through perturbations of either the movement or the feedback signals. The adaptive learning and after-effects that are observed in neurologically healthy subjects minimize performance errors and/or sensory prediction errors, and are typically considered to reflect an updating of the internal forward model, the internal inverse model (control policy), or both (e.g., Flanagan et al., 2003; Mazzoni & Krakauer, 2006; Taylor et al., 2014).

Within this framework, our model of stuttering proposed that individuals who stutter may have difficulty learning accurate internal models of the complex transformations from neuromuscular activation to vocal tract configurations and from those articulatory movements and postures to acoustic speech output (Max, 2004; Max et al., 2004; Neilson & Neilson, 1987). Developmental stuttering has its onset during early childhood when substantial anatomical changes take place in the CNS as well as in the peripheral structures of the vocal tract and, consequently, also in the resulting speech kinematics and acoustics (Callan et al., 2000; Kent, 1976; Kent, 1997; Vorperian et al., 1999; Vorperian et al., 2005; Vorperian & Kent, 2007; Vorperian et al., 2009). If individuals who stutter are impaired in their ability to update internal models of these intricate relationships, it would be problematic for the inverse model to derive accurate motor commands for achieving the desired sensory outcomes and for the forward model to precisely predict the sensory consequences of the planned motor commands. Hypothetically, the resulting sensory prediction errors could trigger maladaptive system corrections that interfere with fluent speech production.

Consistent with this theoretical perspective, recent studies have found that adults who stutter (AWS) exhibit reduced auditory-motor adaptation as compared with adults who do not stutter (AWNS) (Daliri et al., 2018; Daliri & Max, 2018; Sengupta et al., 2016). However, it remains controversial whether such poor sensorimotor learning can be directly related to the mechanisms that are responsible for stuttering given that Daliri et al. (2018) reported that adaptation limitations are not present in *children* who stutter (CWS). Hence, those authors argued that the learning limitations found in AWS reflect a compensatory effect that arises as a result of years of experiencing stuttering.

A second controversy relates to the question whether sensorimotor learning problems in individuals who stutter are specific to the speech system. Differences in movement *control* between stuttering and nonstuttering individuals certainly are not limited to speech tasks, and have been found in behavioral measures of finger movement sequencing accuracy, initiation and execution time (Webster, 1997), manual reaction time (Bishop et al., 1991; Webster & Ryan, 1991), bimanual coordination (Forster & Webster, 2001; Zelaznik et al., 1997), manual responses to visual feedback (Jones et al., 2002), and directional and target accuracy of reaching movements (Daliri et al., 2014). With regard to motor *learning*, several studies have found finger-tapping sequence learning difficulties in AWS (Smits-Bandstra, De Nil et al., 2006a; Smits-Bandstra, De Nil et al., 2006b; Smits-Bandstra & De Nil, 2007; Smits-Bandstra & De Nil, 2009). Yet, the process of nonspeech sensorimotor adaptation in response to feedback perturbations has not been explored in stuttering. This is surprising because the upper-limb sensorimotor learning literature offers various paradigms that have been extensively tested and that are well established. One common paradigm investigates sensorimotor learning by experimentally applying a visuomotor rotation such that the location of a cursor representing hand position is rotated around the center of the workspace during reaching movements (while the actual hand remains invisible). Participants adapt by reaching in the direction opposite to that induced by the visual feedback (e.g., Krakauer & Mazzoni, 2011; Mazzoni & Krakauer, 2006; Shadmehr et al., 2010).

The studies presented here were designed to directly investigate both speech auditory-motor learning and reach visuomotor learning in AWS as well as CWS. We tested age-, sex-, and handedness-matched stuttering and nonstuttering adults and children in each of the two paradigms. Experiment 1 (*N* = 60) involved a speech auditory-motor adaptation task in which formant frequencies in the real-time auditory feedback were shifted up or down either gradually over many trials or suddenly between two successive trials. Experiment 2 (*N* = 84) involved a reach visuomotor adaptation task in which real-time visual feedback was suddenly rotated counterclockwise around the center of the workspace.

## EXPERIMENT 1: SPEECH AUDITORY-MOTOR ADAPTATION IN ADULTS AND CHILDREN WHO STUTTER

### Method

#### Adult participants

We recruited 18 AWS and 18 age- (± 3 years), sex-, and handedness-matched AWNS who reported not taking any medications with a possible effect on sensorimotor functioning and no diagnosed neurological, psychological, emotional, or speech-language-hearing problems (other than stuttering in case of AWS). Each stuttering participant was individually matched with a nonstuttering control participant. We included only native speakers of American English (or individuals who were bilingual but had started speaking English before the age of 5 years). In addition, for AWS, the onset of stuttering had to have occurred during childhood (age 8 years or younger). All participants provided written informed consent in accordance with procedures approved by a local Institutional Review Board.

An American Speech-Language-Hearing Association certified speech-language pathologist administered the Stuttering Severity Instrument (SSI-3 or SSI-4, Riley, 1994; Riley, 2009) to all participants. Based on these SSI results, we excluded three AWS because their speech samples failed to reach an overall SSI score of 10, the minimum score that is required for adults to reach the lowest possible severity classification of “very mild.” On the other hand, one additional AWS was excluded due to being unable to produce a sufficient number of fluent trials during the actual speech adaptation task. Consequently, the final data set included 14 AWS (age *M* = 24.9 years, *SD* = 7.8 years, range 18-49 years) and 14 AWNS (*M* = 25.0 years, *SD* = 7.9 years, range 19-48 years).

Stuttering severity for the remaining AWS ranged from very mild to severe (3 very mild, 3 mild, 5 moderate, 3 very severe). In each group, there were 10 men and 4 women. The groups were matched to include 12 right-handed and 2 left-handed individuals (per self-report). Pure tone air-conduction hearing screenings were completed for all octave frequencies from 250-4000 Hz. Most of the participants had thresholds at or below 20 dB HL for all tested frequencies. One stuttering participant had a threshold of 40 dB HL at 4 kHz for the left ear whereas another stuttering participant also had a threshold of 40 dB HL at 4 kHz but for the right ear; one other stuttering participant had a threshold of 25 dB HL at 4 kHz for the left ear.

#### Child participants

Data were successfully collected for 16 CWS and 16 individually age- (± 3 months)-, sex-, and handedness-matched children who do not stutter (CWNS). A guardian or parent of each child provided written informed consent following procedures approved by a local Institutional Review Board. The children were further divided into two subgroups based on age: 8 CWS and 8 CWNS in a younger age group from 3-6 years old (CWS: *M* = 5.41 years, *SD* = 1.36 years, range = 3.50-6.83 years; CWNS: *M* = 5.49 years, *SD* = 1.29 years, range = 3.75-6.83 years), and 8 CWS and 8 CWNS in an older age group from 7-9 years old (CWS: *M* = 8.02 years, *SD* = 0.69 years, range 7.08-9.33 years; CWNS: *M* = 8.05 years, *SD* = 0.60 years, range = 7.25-9.33 years). Generally consistent with early childhood changes in the distribution of affected girls versus boys, the younger age groups each included 3 girls and 5 boys whereas the older age groups included only boys.

Children’s handedness was determined with a custom-modified version of the Edinburgh Handedness Inventory (Coren, 1993; Oldfield, 1971). Each child was asked to perform 5 tasks (drawing, throwing, using children’s scissors, using a spoon, opening the lid of a box) selected from the 10-items normally included in the Edinburgh inventory. In the younger age groups, seven CWS were right-handed (defined here as completing 4 or 5 of the tasks with the right hand) and one was ambidextrous (defined here as completing 2 or 3 items with the right hand and the other 3 or 2 items with the left hand) whereas all CWNS were right-handed. In the older age groups, all CWS and CWNS were right-handed, based on the same criteria.

All child participants (a) spoke American English as their native language, (b) had normal pure tone air-conduction hearing thresholds (20 dB HL or better for all octave frequencies from 250-4000 Hz in both ears), and (c) scored above the 20^th^ percentile on each of the following speech and language tests: Peabody Picture Vocabulary Test (PPVT-4, Dunn & Dunn, 2007), Expressive Vocabulary Test (EVT-2, Williams, 2007), Goldman-Fristoe Test of Articulation (GFTA-2, Goldman & Fristoe, 2000; GFTA-3, Goldman & Fristoe, 2015), and either Test of Early Language Development (TELD-3, Hresko et al., 1999) or (for children 8 or 9 years old) Clinical Evaluation of Language Fundamentals (CELF-4, Semel et al., 2004). Specific eligibility criteria for children who stutter included (a) confirmation of the diagnosis of stuttering by an American Speech-Language-Hearing Association certified speech-language pathologist, (b) a Stuttering Severity Instrument (SSI-3 or SSI-4, Riley, 1994; Riley, 2009) score of at least 6 (i.e., the minimum score required for children to reach the lowest possible severity classification of “very mild”), and (c) stuttering onset prior to age 5. Based on SSI scores, the stuttering severity classification was severe for 1 child, moderate for 7 children, mild for 5 children, and very mild for 3 children.

#### Experimental set-up

All participants completed the speech adaptation task in a large sound booth. For each trial, adults spoke one of three monosyllabic words (“*tech*,” *“tuck*,” “*talk*”) into a microphone (SM58, Shure). The words were presented on a computer monitor in randomized order per block of three trials. Children named pictures that were used in a board game: each trial elicited the production of a monosyllabic word (*“buck”, “bus”, “puck”, “pup”, “cut”, “cup”, “gut”, “duck”*) while their speech was recorded with a wireless lapel microphone (WL185 with transmitter ULX1-M1 and receiver ULXP4, Shure). The pictures were presented in random order per block of 8 trials.

As illustrated in Figure 1A, the microphone signal was amplified (DI/O Preamp System II, ART) and recorded on one channel of a CD recorder (CD-RW901SL, Tascam). The same signal was also routed to a vocal processor (VoiceOne, TC Helicon) that can manipulate all formant frequencies (i.e., resonance frequencies of the vocal tract) in real-time with a total feedback loop delay of only 10 ms^1^ (Kim et al., 2020). The processor’s output was routed to a headphones amplifier (S-phone, Samson) and insert earphones (ER-3A, Etymotic Research) worn by the participant. Before each participant’s recording session, this feedback loop was calibrated such that speech with an intensity of 75 dB SPL at the microphone (15 cm from a loudspeaker through which a production of “tuck” was played) resulted in an output of 72 dB SPL in the earphones (measured in a 2 cc coupler Type 4946 connected to a sound level meter Type 2250A with Type 4947 1⁄2” pressure field microphone, Bruel & Kjaer). This input-output ratio is based on simultaneous recordings of a speech signal at a microphone in front of a speaker and at the entrance to the speaker’s ear (Cornelisse et al., 1991).

**Figure 1.**
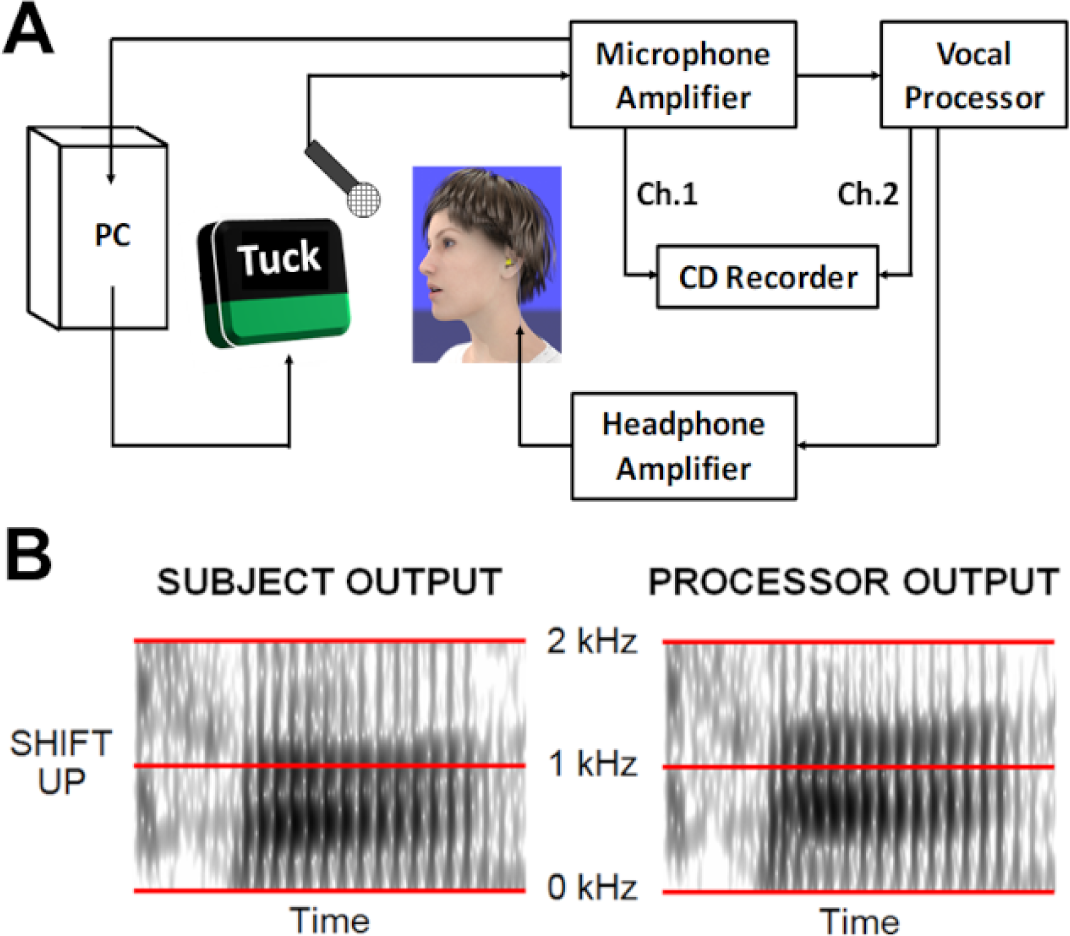
Experimental set-up for speech auditory-motor adaptation. (A) The speech signal from the microphone is routed to a vocal processor that applies a formant shift perturbation to the signal that is presented through insert earphones as real-time auditory feedback. (B) Effect of an upward formant shift illustrated with separate spectrographic displays of the original acoustic speech signal produced by the participant (left) and the altered signal that is output by the vocal processor and heard by the subject (right).

#### Speech auditory-motor adaptation task

Each session started with a baseline phase without formant perturbation. After the baseline phase, all formant frequencies in the speech signal were shifted either up or down by the VoiceOne processor which was controlled by a computer with custom MATLAB (The Mathworks) code. The processor selectively shifted the formant frequencies but not the fundamental frequency or consonant-related noise components (Figure 1B). We opted to have all formant frequencies shifted by the same proportional amount and in the same direction. This manipulation does not induce the perception of phonemic errors (a manipulation that we applied in Feng et al., 2011), but implements a different motor-to-sensory transformation corresponding to speech output from a vocal tract with increased or decreased geometrical dimensions (as we have also applied in Daliri & Max, 2018; Max et al., 2003; Max & Maffett, 2015). Thus, a participant’s articulatory movements resulted in real-time auditory feedback with resonance frequencies that were globally increased or decreased relative to those associated with the participant’s own motor-to-sensory mapping.

For adults, the processor implemented upward or downward formant shifts of 250 cents (the difference in cents between two frequencies fa and fb is 1200 × log2(fa/fb)). For each of the two shift directions there was one condition in which the formant perturbation was introduced suddenly (i.e., the maximum perturbation was introduced in full as a single step between two successive trials) and one condition in which the formant perturbation was introduced gradually (i.e., incrementally ramped up or down to its maximum value across many trials). Thus, there were four conditions: sudden shift up, gradual shift up, sudden shift down, and gradual shift down. For children, to make the perturbation sufficiently salient and limit the testing duration, we selected a shift of 335 cents but only in the upward direction. Thus, children completed two conditions: sudden shift up and gradual shift up.

The decision to include both sudden (or step) and gradual (or ramp) introductions of the auditory perturbation was made based on others’ suggestions that intact cerebellar functioning is critical specifically for adapting to gradual perturbations (Doya, 2000; Robertson & Miall, 1999) whereas intact basal ganglia functioning is critical specifically for adapting to sudden perturbations (Contreras-Vidal & Buch, 2003; Doya, 2000; Venkatakrishnan et al., 2011)— suggestions that were deemed potentially informative for the present study given that dysfunction of both these neural substrates has been implicated in stuttering (e.g., Civier et al., 2013; Ingham et al., 2004). It should be noted, however, that the relationship between neural substrates and adaptation is complex and poorly understood: other empirical results suggest that patients with basal ganglia disease can be unimpaired when adapting to sudden visuomotor distortions or mechanical force fields during reaching (Gutierrez-Garralda et al., 2013; Smith & Shadmehr, 2005; Weiner et al., 1983) whereas cerebellar patients may be (a) impaired in adapting to sudden visuomotor or force field perturbations during reaching (e.g., Criscimagna-Hemminger et al., 2010; Maschke et al., 2004; Rabe et al., 2009; Schlerf et al., 2013; Smith & Shadmehr, 2005; Weiner et al., 1983; Werner et al., 2009; Werner et al., 2010) and (b) relatively unimpaired when adapting to a gradual force field perturbations during such reaching movements (e.g., Criscimagna-Hemminger et al., 2010). Thus, the relationship between cortico-striato-thalamo-cortical or cortico-cerebellar-thalamo-cortical circuits and impaired adaptation to sudden versus gradual perturbations is clearly not as straightforward as had been suggested. In addition, there is also evidence that both neural circuits may contribute to different stages of the same adaptation process (Doyon et al., 2003) and that the cerebellum may process performance-related information (e.g., an error signal) that is a pre-requisite for adaptation but that is not directly related to learning *per se* (Werner et al., 2009).

The order of completion of the conditions was pseudo-balanced (or balanced in children) across stuttering individuals, and each nonstuttering individual performed the four conditions (or two conditions for children) in the same order that had been used for the matched stuttering participant. Adult participants completed for each condition 60 blocks of the three test words (180 words) at a rate of 5 blocks (15 words) per minute. In the adults’ gradual conditions, there were 10 blocks of *baseline*, 20 blocks of *gradual ramp*, 20 blocks of *full exposure* (i.e., 250 cents up or down formant shift), and 10 blocks of *after-effects* or wash-out (i.e., veridical feedback was restored). In the adults’ sudden conditions, there were 20 blocks of *baseline*, 30 blocks of *full exposure*, and 10 blocks of *after-effects*. Intensity of the subject’s speech was kept relatively constant across trials by means of color-coded visual feedback on the computer monitor. Words were presented on the monitor at a consistent pace so that subjects read 15 words per minute.

Children completed for each condition 13 blocks of the eight test words. Even though the use of a board game prevented us from controlling the rate of trial presentation as strictly as we did for adults, the staff member interacting with the children always attempted to keep the pace at approximately 10-12 trials per minute. In the children’s gradual condition, there were 2 blocks of *baseline*, 4 blocks of *gradual ramp*, 5 blocks of *full exposure* (i.e., 335 cents upward formant shift), and 2 blocks of *after-effects*. In their sudden condition, there were 4 blocks of *baseline*, 7 blocks of *full exposure*, and 2 blocks of *after-effects*.

#### Data extraction and analyses

Prior to analyzing formant frequency changes to quantify auditory-motor adaptation, median vowel duration was measured for each participant in each condition. This analysis served to rule out the possibility that one group benefited from a slower rate of speech which would allow more time for *within-trial* compensation based on online sensory feedback (as opposed to sensorimotor adaptation which affects moving planning based on feedback from previous trials).

The acoustic recordings were then resampled at 10 kHz, and the frequencies of the first (F1) and second (F2) formant were measured in the middle of the vowel using a custom MATLAB program. The window defining vowel midpoint had its onset 40% into the vowel and its offset 60% into the vowel (note that for 96.05% of all trials from the adult participants and 97.90% of all trials from the child participants, this analysis window did not extend beyond 150 ms after vowel onset; thus, any effects of online feedback-driven compensation can be expected to be minimal). Within the MATLAB program, we extracted children’s formants using custom code implementing an order-optimized Linear Predictive Coding method (Feng et al., 2011; Vallabha & Tuller, 2002) and adults’ formants using calls to Praat scripts (Boersma & Weenink, 2008). F1 and F2 in each trial were converted to cents normalized to the participant’s baseline values (calculated from blocks 6-10 for adults and block 2 for children). F1 and F2 were then averaged for all analyses reported here. Trials in which stuttering occurred, the word was mispronounced, or pronunciation was affected by yawning, etc., were excluded (0.6% of all trials for adults, 3.0% of all trials for children).

Both the final extent of adaptation and initial or early adaptation were determined for each participant in each condition. Measures of the final extent of adaptation were based on the last five blocks from the full perturbation phase for adults and the last two blocks from the full perturbation phase for children. Measures of initial adaptation were based on early trials in the full perturbation phase of the conditions with sudden onset of the perturbation, specifically the initial five blocks for adults and the first block for children. We were not able to estimate initial *rate* of learning by fitting exponential functions adaptation because the auditory-motor adaptation profiles of many AWS and CWS did not demonstrate exponential learning.

Statistical analyses were completed in R (R Core Team, 2018). Using the *aov* function, we conducted analyses of variance (ANOVA) separately for final extent of adaptation and initial adaptation. For adults, we used repeated measures ANOVA to test the between-groups effect (stuttering vs. nonstuttering) as well as within-groups effects related to perturbation type (sudden vs. gradual) and perturbation direction (i.e., up-shift vs. down-shift) as well as all interactions among these variables.^2^ For children, who completed only up-shift conditions, the ANOVA included two between-group factors (stuttering vs. nonstuttering and 3-6 vs. 7-9 years of age).

The analysis of children’s final extent of adaptation was a repeated measures ANOVA that also included the within-group effect of perturbation type (sudden vs. gradual); the analysis of children’s initial adaptation included no within-group variable as this measure was extracted only for the sudden perturbation condition. Effect sizes were calculated as partial omega-squared (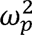) using the *MOTE* package (Buchanan et al., 2019). Welch’s *t*-tests were used in all cases where ANOVA results necessitated follow-up with pair-wise comparisons. In those instances, we report *p* values adjusted with the Holm-Bonferroni method (Holm, 1979) to keep family-wise error rates at .05. When between-group differences were identified, one-sample *t*-tests with Holm-Bonferroni corrections were used to determine which groups or sub-groups did or did not adapt (i.e., showed significant change from baseline). Lastly, we conducted two sets of correlational analyses with Pearson’s correlation coefficient. First, for the stuttering adults and children, we examined whether there was a relationship between either final extent of adaptation or initial adaptation and stuttering frequency (averaged percent stuttered syllables across reading and speaking samples from the SSI). Second, for stuttering and nonstuttering children, we examined whether there was a relationship between either final extent of adaptation or initial adaptation and age.

## Results

### Vowel duration

Although AWS tended to produce longer vowels than AWNS during the auditory-motor learning task (*M* = 152 ms, *SD* = 74 ms, and *M* = 112 ms, *SD* = 21 ms, respectively), the difference was not statistically significant, *F*(1, 26) = 3.806, *p* = 0.062, 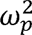 = 0.049. Vowel duration also did not differ between the conditions with upward vs. downward directions of formant perturbation, *F*(1, 26) = 0.496, *p* = 0.487, 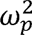 = 0.000, or with sudden vs. gradual types of perturbation, *F*(1, 26) = 3.491, *p* = 0.073, 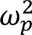 = 0.001). None of the interactions were statistically significant (Direction × Group: *F*(1, 26) = 0.960, *p* = 0.336, 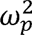 = 0.000; Type × Group: *F*(1, 26) = 0.355, *p* = 0.557, 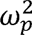 = 0.000; Direction × Type: *F*(1, 26) = 0.533, *p* = 0.472, 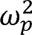 = 0.000; Direction × Type × Group: *F*(1, 26) = 0.002, *p* = 0.965,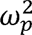 = -0.001).

Similarly, measures of vowel duration for CWS and CWNS (*M* = 110 ms, *SD* = 40 ms, and *M* = 117 ms, *SD* = 32 ms, respectively) were not statistically significantly different, *F*(1, 28) = 0.269, *p* = 0.608, 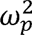 = -0.013. There was also no statistically significant difference in vowel duration between the two age-based subgroups of children, *F*(1, 28) = 0.083, *p* = 0.775, 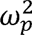 = -0.016, or between the two types of perturbation, *F*(1, 28) = 2.800, *p* = 0.105, 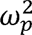 = 0.006. Finally, none of the interactions were statistically significant (Age × Group, *F*(1, 28) = 0.416, *p* = 0.524, 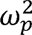 = -0.001; Type × Group, *F*(1, 28) = 1.289, *p* = 0.266,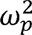 = 0.001; Age × Type, *F*(1, 28) = 0.007, *p* = 0.932,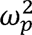= -0.003; Group × Age × Type, *F*(1, 28) = 0.209, *p* = 0.651,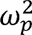 = -0.003).

### Auditory-motor adaptation in AWS vs. AWNS

Figure 2 shows adult group data (means and standard errors of the mean) across all trials in the sudden and gradual up-shift conditions. Formant frequencies are represented in cents relative to baseline (i.e., average frequency for the blocks from which baseline is calculated is always 0 cents). Participants lowered their formant frequencies during the perturbation trials in both the sudden and gradual shift conditions. Figure 3 shows the corresponding data for the down-shift conditions. In this case, participants tended to increase their formant frequencies, albeit to a smaller extent. The ANOVA results for final extent of auditory-motor learning (per participant averaged across F1 and F2, the 3 test words, and trial blocks 46-50) revealed statistically significant main effects for Group, *F*(1, 26) = 9.780, *p* = 0.004, 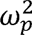 = 0.140, and Direction, *F*(1,26) = 20.738, *p* < 0.001, 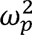 = 0.195: AWS adapted significantly more than AWNS, and upward formant shift perturbations resulted in significantly more adaptation than downward formant shift perturbations. The main effect for perturbation Type was not significant, *F*(1, 26) = 0.719, *p* = 0.404, 0.040, 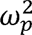 = -0.002, but there was a significant Group × Type interaction, *F*(1, 26) = 4.696, *p* = 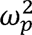 = 0.021. This interaction was due to the AWS vs. AWNS between-group difference in final extent of adaptation being larger for the sudden perturbation than for the gradual perturbation. Nevertheless, post-hoc tests revealed that the between-group difference was statistically significant in both the sudden perturbation condition, *t*(19.135) = -3.337, *p* = 0.007, *d* = 1.261, and the gradual perturbation condition *t*(20.687) = -2.298, *p* = 0.032, *d* = 0.869. There were no other statistically significant interactions for the final extent of adaptation (Direction × Group: *F*(1, 26) = 0.243, *p* = 0.626, 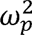 = -0.009; Type × Direction: *F*(1, 26) = 0.282, *p* = 0.600, 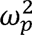 = -0.003; Type × Direction × Group: *F*(1, 26) = 0.177, *p* = 0.678, 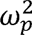 = -0.004). A family of one-sample *t*-tests demonstrated that the final extent of adaptation for AWNS was statistically significantly different from baseline in all four conditions (sudden up-shift, *t*(13) = -9.056, *p* < 0.001, *d* = 2.420, gradual up-shift, *t*(13) = -10.881, *p* < 0.001, *d* = 2.908, sudden down-shift, *t*(13) = 4.920, *p* = 0.001, *d* = 1.315, gradual down-shift, *t*(13) = 3.060, *p* = 0.009, *d* = 0.818). In contrast, AWS showed adaptation only in the gradual up-shift condition (*t*(13) = - 4.472, *p* = 0.003, *d* = 1.195) but not in the other three conditions (sudden up-shift, *t*(13) = -2.089, *p* = 0.171, *d* = 0.558, sudden down-shift, *t*(13) = -0.288, *p* = 1.000, *d* = 0.077, gradual down-shift, *t*(14) = 0.473, *p* = 1.000, *d* = 0.126).

**Figure 2.**
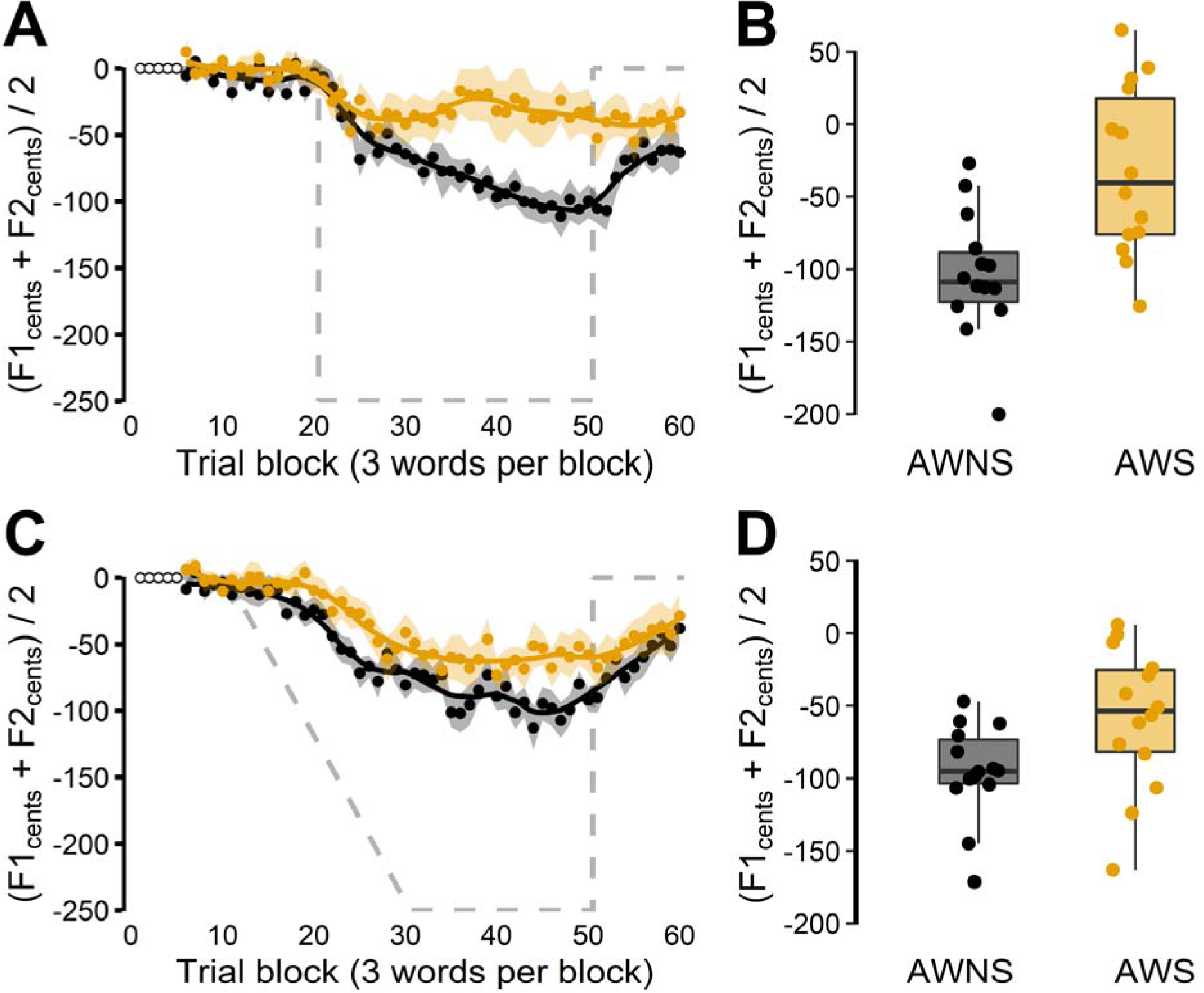
Auditory-motor adaptation for adults who do not stutter (black) and adults who stutter (orange) in the 250 cents upward formant perturbation conditions. Panels A (suddenly introduced perturbation) and C (gradually introduced perturbation) show group averaged formant frequencies in cents relative to baseline (averaged across F1 and F2 and across 3 test words after normalization of the individual formants and words). Shaded areas indicate standard errors of the mean. Gray dashed lines plot the formant perturbation with reversed sign (thus indicating hypothetical full compensation). Panels B and D show corresponding individual participant data (averaged across the last 5 blocks of 3 words) overlaid on group boxplots.

**Figure 3.**
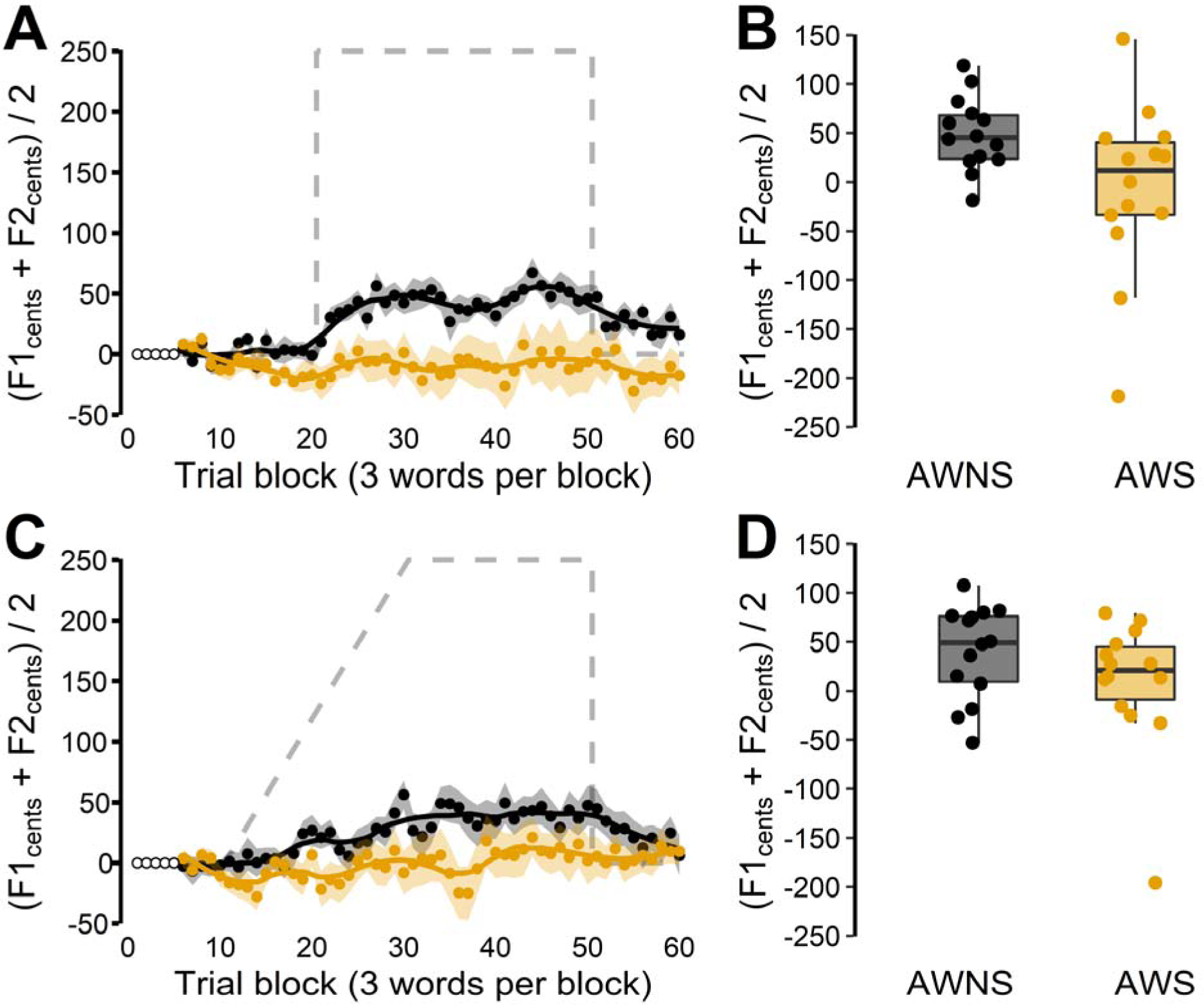
Auditory-motor adaptation for adults who do not stutter (black) and adults who stutter (orange) in the 250 cents downward formant perturbation conditions. Panels A (suddenly introduced perturbation) and C (gradually introduced perturbation) show group averaged formant frequencies in cents relative to baseline (averaged across F1 and F2 and across 3 test words after normalization of the individual formants and words). Shaded areas indicate standard errors of the mean. Gray dashed lines plot the formant perturbation with reversed sign (thus indicating hypothetical full compensation). Panels B and D show corresponding individual participant data (averaged across the last 5 blocks of 3 words) overlaid on group boxplots.

We examined adults’ initial adaptation using the first five perturbation blocks of the sudden conditions. ANOVA revealed a significant between-group difference, *F*(1, 26) = 6.267, *p* = 0.019, 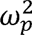 = 0.089, with greater initial adaptation for AWNS than for AWS. There was also a significant main effect of perturbation direction, *F*(1, 26) = 4.872, *p* = 0.036, 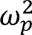 = 0.057, with greater initial learning for upward perturbations than for downward perturbations. However, these two main effects were modified by a Group × Direction interaction, *F*(1, 26) = 4.352, *p* = 0.047, 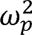 = 0.049. Post hoc tests revealed that AWS, as compared with AWNS, learned less in the initial blocks of the sudden down condition, *t*(19.701) = -3.483, *p* = 0.005, *d* = 1.317, but not in the sudden up condition, *t*(22.423) = -0.527, *p* = 0.604, *d* = 0.199.

There was no statistically significant correlation between stuttering frequency (averaged across the reading and speaking samples from the SSI evaluation) and either final adaptation extent (Sudden, *r*(12) = -0.041, *p* = 0.888, Gradual, *r*(12) = 0.171, *p* = 0.560) or initial adaptation (*r*(12) = 0.211, *p* = 0.470).

### Auditory-motor adaptation in CWS vs. CWNS

In both the sudden and gradual perturbation conditions, CWNS showed a final extent of auditory-motor adaptation that was very similar to that seen in nonstuttering adults (Figure 4 for the younger subgroup 3-6 years of age, Figure 5 for the older subgroup 7-9 years of age). Even the youngest subgroup of CWNS adapted in both conditions to an extent that compensated for 30∼40% of the perturbation, which is highly similar to the data described above for nonstuttering adults. On the other hand, both younger and older CWS made only minimal, if any, changes in their formant frequencies in response to the perturbation. In fact, based on a family of four one-sample *t*-tests, neither the younger nor the older subgroup of CWS showed a final extent of adaptation (per participant averaged across F1 and F2, the different tests words, and blocks 10 and 11) that was significantly different from baseline in either of the two perturbation conditions (Age 3-6 in the sudden condition, *t*(7) = 0.611, *p* = 1.000, *d* = 0.216; Age 3-6 in the gradual condition, *t*(7) = 0.135, *p* = 1.000, *d =* 0.048; Age 7-9 in the sudden condition, *t*(7) = -2.175, *p =* 0.198, *d* = 0.769; Age 7-9 in the gradual condition, *t*(7) = -2.411, *p* = 0.187, *d* = 0.852. On the other hand, the family of one-sample *t*-tests for CWNS indicated that final adaptation of both the younger and the older subgroups differed significantly from baseline in both perturbation conditions (Age 3-6 in the sudden condition, *t*(7) = -6.184, *p* = 0.002, *d* = 2.186, Age 7-9 in the gradual condition, *t*(7) = -4.529, *p* = 0.005, *d* = 1.601, Age 7-9 in the sudden condition, *t*(7) = - 3.788, *p* = 0.007, *d* = 1.339, Age 7-9 in the gradual condition, *t*(7) = -5.172, *p* = 0.004, *d* = 1.829).

**Figure 4.**
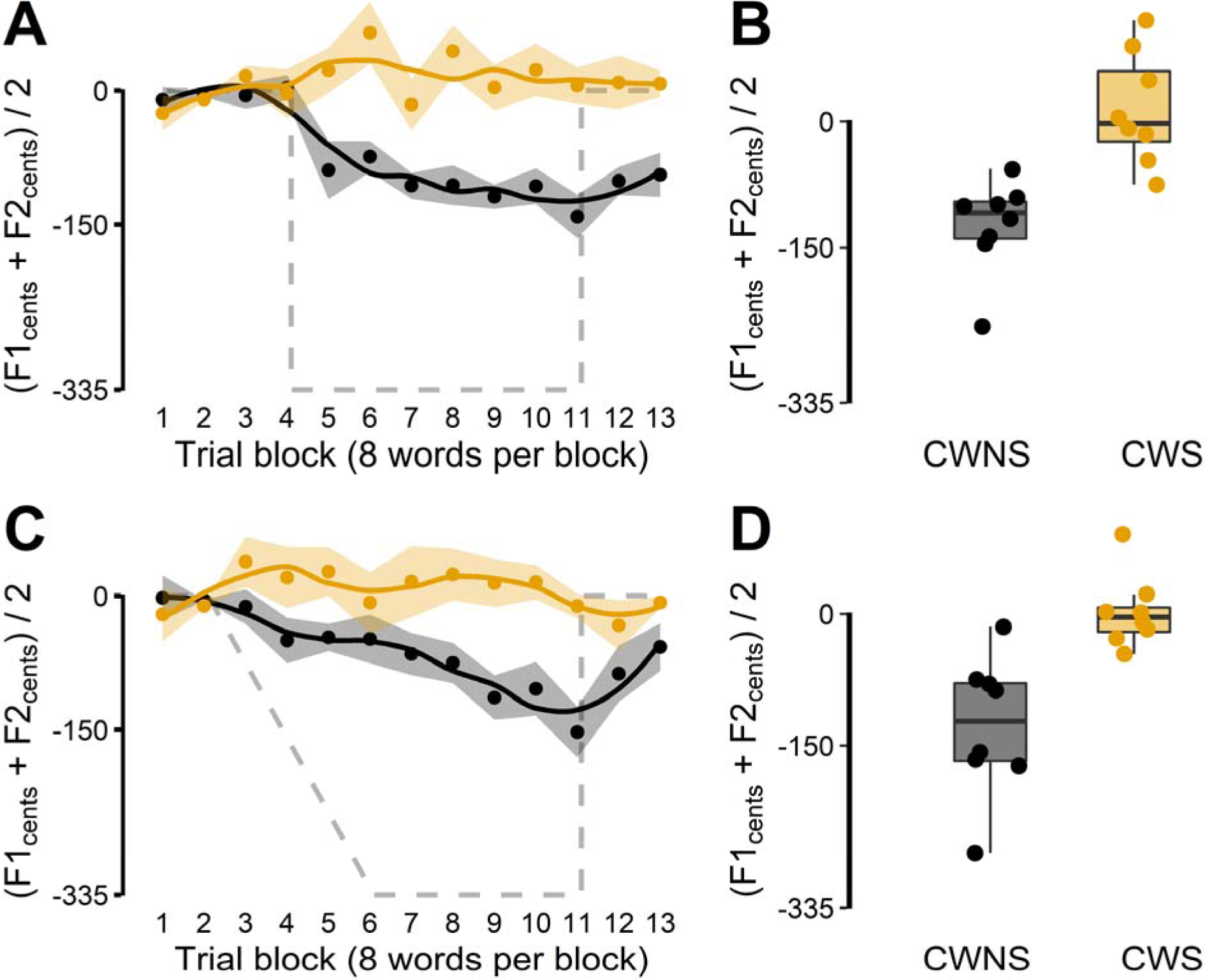
Auditory-motor adaptation for 3-6 year-old children who do not stutter (black) and age-matched children who stutter (orange) in 335 cents upward formant perturbation conditions. Panels A (suddenly introduced perturbation) and C (gradually introduced perturbation) show group averaged formant frequencies in cents relative to baseline (averaged across F1 and F2 and across 8 test words after normalization of the individual formants). Shaded areas indicate standard errors of the mean. Gray dashed lines plot the formant perturbation with reversed sign (thus indicating hypothetical full compensation). Panels B and D show corresponding individual participant data (averaged across the last 2 blocks of 8 words) overlaid on group boxplots.

**Figure 5.**
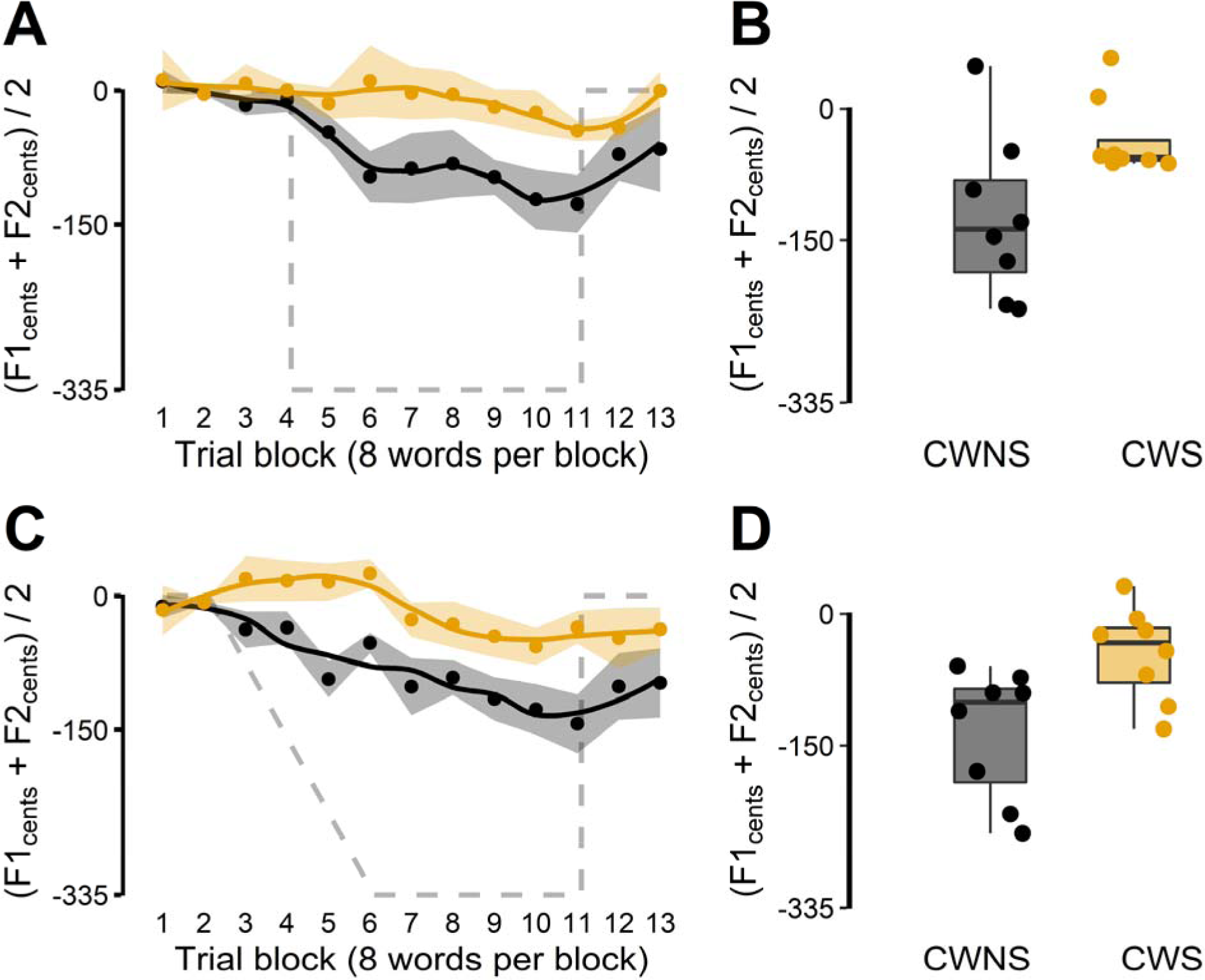
Auditory-motor adaptation for 7-9 year-old children who do not stutter (black) and age-matched children who stutter (orange) in 335 cents upward formant perturbation conditions. Panels A (suddenly introduced perturbation) and C (gradually introduced perturbation) show group averaged formant frequencies in cents relative to baseline (averaged across F1 and F2 and across 8 test words after normalization of the individual formants). Shaded areas indicate standard errors of the mean. Gray dashed lines plot the formant perturbation with reversed sign (thus indicating hypothetical full compensation). Panels B and D show corresponding individual participant data (averaged across the last 2 blocks of 8 words) overlaid on group boxplots.

Consequently, the ANOVA for children’s final extent of adaptation revealed a significant between-group effect, *F*(1, 28) = 41.947, *p* < 0.001, 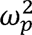 = 0.414, in the absence of main effects for age subgroup, *F*(1, 28) = 2.257, *p* = 0.144, 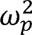 = 0.021, or perturbation type *F*(1, 28) = 0.385, *p* = 0.540, 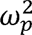 = -0.010. There were also no significant interactions among any of the variables (Age × Group, *F*(1, 28) = 1.684, *p* = 0.205, 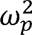 = 0.012; Type × Group, *F*(1, 28) = 0.019, *p* = 0.891, 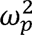 = -0.015; Age × Type, *F*(1, 28) = 0.007, *p* = 0.932, 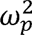 = -0.016; Group × Age × Type, *F*(1, 28) = 0.015, *p* = 0.903, 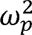 = -0.016).

Despite the absence of a statistically significant Group × Age interaction, inspection of the individual participant data clearly showed that final extent of adaptation increased with age in CWS but not CWNS (Figure 6). In both the sudden and gradual perturbation conditions, the final adaptation extent for CWS was significantly correlated with age, (sudden, *r*(14) = -0.571, *p* = 0.021; gradual, *r*(14) = -0.508, *p* = 0.045). For CWNS, on the other hand, there were no statistically significant correlations in either condition (sudden, *r*(14) = -0.031, *p* = 0.910; gradual, *r*(14) = -0.029, *p* = 0.915). Note that a greater adaptation extent corresponds to a more negative number, and, thus, that a negative correlation here indicates greater adaptation with increasing age.

**Figure 6.**
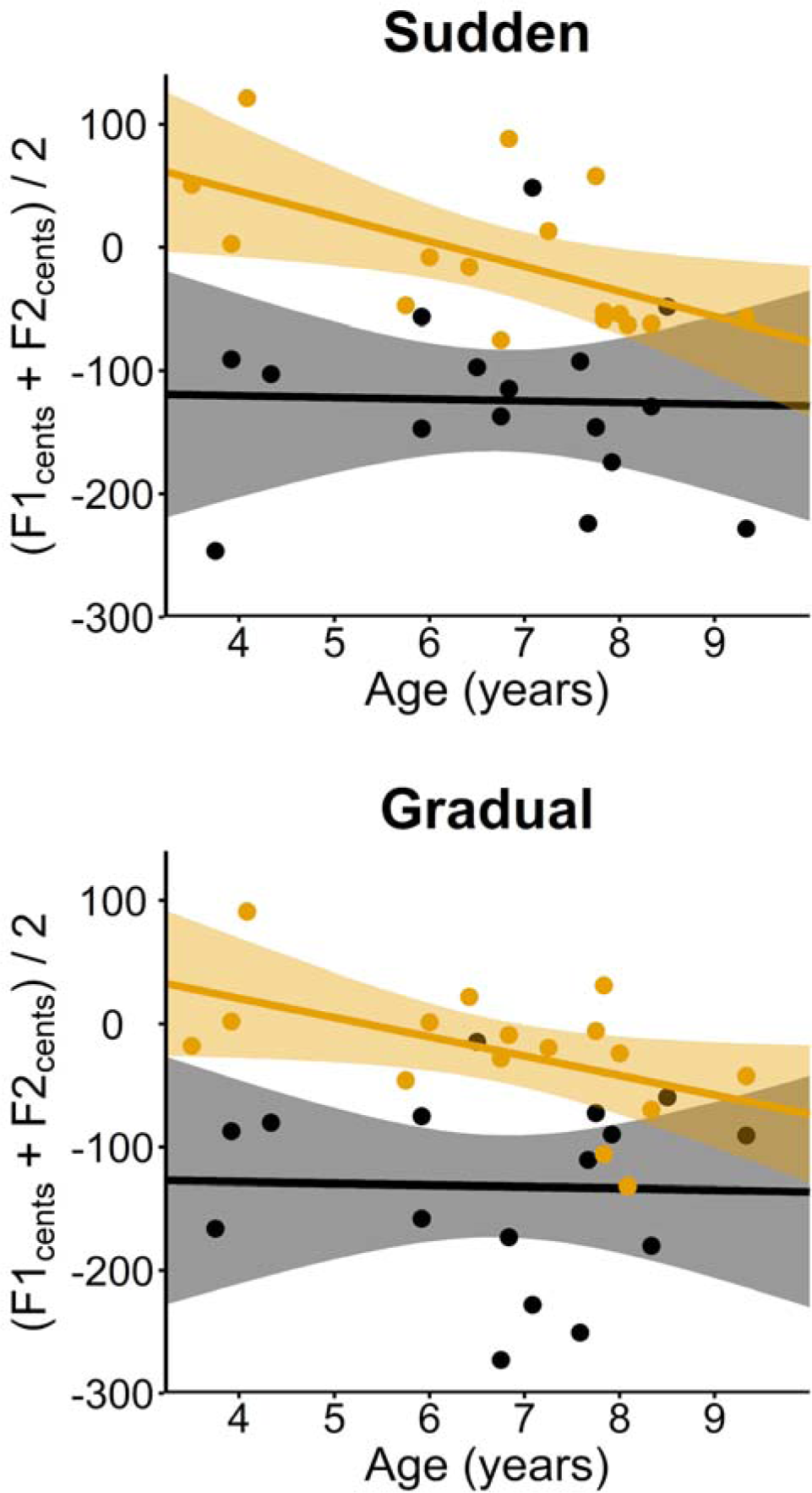
Relationship between extent of speech auditory-motor adaptation and age for CWS (orange) and CWNS (black). Data for the sudden condition shows a negative correlation for CWS whereas there is no such clear trend for CWNS (top). A similar finding in the gradual condition (bottom). The shaded areas indicate the confidence interval (95%).

The ANOVA for children’s initial adaptation in the sudden perturbation condition revealed that CWNS already learned more than matched CWS in the first perturbation block (block 5), *F*(1, 28) = 8.689, *p* = 0.006, 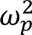 = 0.194. The initial amount of adaptation in this block of trials did not significantly differ between the age subgroups, *F*(1, 28) = 0.012, *p* = 0.913, The Group × Age interaction was also not significant, *F*(1, 28) = 2.642, *p* = 0.115, 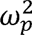 = -0.032. 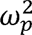 = 0.049.

Lastly, there was no statistically significant correlation between stuttering frequency from the SSI speaking samples and either final extent of adaptation (Sudden, *r*(14) = -0.139, *p* = 0.608, Gradual, *r*(14) = -0.346, *p* = 0.190) or initial adaptation (*r*(14) = 0.266, *p* = 0.319).

## Discussion

In this first experiment, we examined adaptation to formant-shifted real-time auditory feedback in adults who stutter and children who stutter as compared with matched control participants who do not stutter. Specifically, the auditory perturbations consisted of global formant shifts (i.e., applying the same proportional amount of shift to all formants) such that the feedback signal corresponded to vowels produced with a vocal tract that is smaller (in the case of upward shifts) or larger (in the case of downward shifts) than the speaker’s own vocal tract.

These perturbations were applied continually while participants produced *CVC* words, where *C* was always a stop consonant and *V* was a front, central, or back mid vowel. The speaker’s fundamental frequency and consonant bursts were not affected by the perturbation. In other words, the formant shifts specifically changed the transformation from motor commands for vowel-related vocal tract configurations to acoustic output (typically leading the CNS to update its internal models of these transformations), but without changing the phonemic category of the produced vowels or the primary characteristics of the surrounding consonants.

As a first finding, we replicated previous results indicating that AWS show reduced speech auditory-motor learning in response to formant perturbations that are introduced suddenly in-between two successive trials (Daliri & Max, 2018; Sengupta et al., 2016) as well as formant perturbations that are introduced gradually across multiple trials (Daliri et al., 2018). Second, we found that this between-group difference in auditory-motor learning for AWS vs. AWNS holds up for both up-shift and down-shift global formant perturbations. Third, although adaptation for AWS was statistically significantly smaller than that for AWNS in both the suddenly introduced and the gradually introduced perturbation conditions, the between-group difference was larger when the perturbation was introduced suddenly. Fourth, not only did we find a difference in speech auditory-motor learning also in CWS vs. CWNS, the effect size associated with the between-group difference in final extent of adaptation for children (41.4% of variance accounted for) was almost three times as large as that found for adults (14.0% of variance accounted for).

The youngest group of CWS (3-6 years of age) showed essentially no learning at all in either the sudden or the gradual perturbation condition whereas the age-matched CWNS already showed adult-like learning. Descriptively, the older group of CWS (7-9 years of age) showed slightly more adaptation, but even for this group the change from baseline was still not statistically significant for either perturbation type condition. Fifth, like the situation for adults, stuttering and nonstuttering children differed in adaptation for both suddenly and gradually introduced formant perturbations. For children, however, this between-group difference was not affected by the sudden or gradual manner in which the perturbation was introduced as, statistically, there was no Group by Condition interaction. Sixth, among CWS, the final extent of speech auditory-motor learning was correlated with age (more adaptation in older children); in contrast, for CWNS there was no correlation between these variables as even the younger children already adapted to the same extent as older children (and even as adults, but formant perturbation size as well as sample size differed between the child and adult groups so no direct statistical comparisons were made for children vs. adults).

Clearly, our findings that (a) speech auditory-motor learning problems are present not only in AWS but also in CWS, (b) this learning impairment in CWS is profound, and (c) CWS’ learning impairment decreases in magnitude with age, are not consistent with previous work by Daliri et al. (2018). Those authors reported that speech auditory-motor learning limitations occur only in adults who stutter (who have experienced stuttering for many years and whose performance may be affected by compensatory sensorimotor strategies) and not in children who stutter (who are closer to the onset of the disorder). Thus, our contrasting findings with regard to children constitute some of the most important outcomes of the present work because for any sensorimotor learning impairment to potentially play a role in the onset of stuttering (which typically occurs between 2 and 4 years of age), the impairment must of course already be influencing the development of speech motor behavior during early childhood.

One possible reason for the discrepancy between our own results and those of Daliri et al. (2018) may lie in the specific formant perturbations that were used in the studies. In our work described here, we shifted all formants in the same direction such that the auditory feedback signal reflected an altered motor-to-sensory transformation that did not change the phonemic category of the produced vowels. In other words, when the children participating in our study produced the test words *bus, pup, cup, duck*, etc., they never perceived these words as having been produced with the wrong vowel. Instead, they perceived the words as having the correct vowel but produced by a vocal tract with different geometric properties (akin to developmental changes in vocal tract geometry during childhood, albeit on a short time scale of only minutes). Daliri et al. (2018), on the other hand, shifted the first and second formants in different directions such that when participating children produced the words *bed, head,* and *Ted,* they heard their productions as similar to *bad, had,* and *Tad*. It is very well possible, and perhaps even expected, that CWS are indeed able to correct their productions when auditory feedback from previous trials indicates that they produced the words with a completely wrong sound (as in Daliri et al., 2018). For example, the fact that CWS are not more likely than other children to have speech sound disorders (Nippold, 2002; Unicomb et al., 2020) suggests that CWS do not have difficulty with adjusting movement planning to target the correct speech sounds. In addition, the symptoms of stuttering do not involve producing incorrect sounds, but certain movements or postures for sound production being repeated or sustained.

It is therefore important to recall that what had been hypothesized previously in a theoretical framework (e.g., Max, 2004; Max et al., 2004) is not that CWS would fail to correct for errors in target sound selection, but, rather, that there may be “an inability to learn stable or correct mappings between motor and sensory signals and to update these mappings in the presence of rapid neural and craniofacial maturation during speech development” (Max, 2004, p. 374). To be more specific, the crux of that particular hypothesis is that “After their initial acquisition during babbling and early speech, these internal models require continuous updating, due to the rapid developmental changes in neural, anatomical, and biomechanical characteristics during childhood. … If, for some reason, the CNS would fail to accurately update the internal models to match the currently applicable transformations, it would become unable to predict with great precision the sensory consequences of planned movements …” (Max, 2004, p. 374). The formant perturbations that we implemented in our work reported here were selected very specifically to test participants’ updating of internal models that represent such fine-grained and vocal tract geometry-related spectral features of the motor-to-auditory transformations involved in speech production. Our findings very strongly suggest that, at least for the time scale tested here, CWS did not update these internal models at all whereas age-matched CWS already showed adult-like learning. Moreover, our correlational analyses demonstrate that younger CWS—who are closer to the onset of stuttering—showed less learning than older CWS, thus rejecting the notion that impaired speech auditory-motor learning is something that develops as a consequence of stuttering rather than a factor that may contribute to the onset of stuttering.

## EXPERIMENT 2: REACH VISUOMOTOR ADAPTATION IN ADULTS AND CHILDREN WHO STUTTER

### Method

#### Adult participants

We recruited 16 AWS, but one participant was excluded due to the SSI score being too low to reach the “very mild” category. Thus, the final data set included 15 AWS (age *M* = 27.07 years, *SD* = 7.88 years, range 19-48 years) and 15 individually age- (±3 years), sex-, and handedness-matched AWNS (age *M* = 26.67 years, *SD* = 8.16 years, range = 20-48 years). All participants had provided written informed consent in accordance with procedures approved the local Institutional Review Board. The same inclusion criteria as in Experiment 1 were applied, except that participants did not need to be native speakers of American English given that the task involved upper limb reaching movements rather than speaking. All participants were right-handed based on self-report, and there were 12 males and 3 females in each group. According to the SSI scores, 2 individuals’ stuttering was categorized as very severe, 1 as severe, 4 as moderate, 4 as mild, and 4 as very mild. Six of the 15 AWS and 4 of the 15 AWNS were also included above in Experiment 1.

#### Child participants

The participating children were 27 CWS and 27 CWNS from 3 to 9 years of age (10 CWS and 2 CWNS also participated in Experiment 1). A guardian or parent provided written informed consent following the procedures approved by a local Institutional Review Board. Inclusion criteria were as described for Experiment 1. Each stuttering child was individually matched with a nonstuttering child based on age (i.e., ±3 months) and sex. Based on the handedness assessment described above for Experiment 1, most children were right-handed except 4 CWS and 1 CWNS who were ambidextrous. We organized the data into three age-based subgroups. A 3-4 years old subgroup included 9 CWS (*M* = 3.80 years, *SD* = 0.57 years, range = 3.11-4.55 years) and 9 CWNS (*M* = 3.76 years, *SD* = 0.52 years, range = 3.26-4.66 years). A 5-6 years old subgroup included 8 CWS (*M* = 6.10 years, *SD* = 0.50 years, range =5.46-6.89 years) and 8 CWNS (*M* = 6.14 years, SD = 0.46 years, min = 5.50, max = 6.69). A 7-9 years old subgroup included 10 CWS (*M* = 8.13 years, *SD* = 0.68 years, range = 7.09-9.13 years) and 10 CWNS (*M* = 8.09 years, *SD* = 0.72 years, range = 7.06-9.09 years). In the 3-4 years-old range, there were 2 girls in each group. In the 5-6 years-old range, there were 3 girls in each group. In the 7-9 years-old range, each group included 1 girl.

With regard to severity based on SSI scores (SSI-3 or SSI-4, Riley, 1994; Riley, 2009), stuttering in the youngest group of CWS (3-4 years of age) was rated as very severe in 1 case, severe in 1 case, moderate in 4 cases, and mild in 3 cases. For the group of CWS in the range 5-6 years old, stuttering was severe for 1 child, moderate for 3 children, mild for 2, and very mild for 2. The oldest group of CWS, 7-9 years of age, included 2 children rated as severe, 4 rated as moderate, 3 rated as mild, and 1 rated as very mild.

#### Experimental set-up

Seated on a height-adjustable chair, participants made fast out-and-back reaching movements with their right arm supported by an acrylic air sled on a glass table top (Figure 7A). The air sled was connected to a compressed-air supply so that friction across the glass surface was minimized. The position of a sensor attached to the participant’s extended right index finger was recorded by an electromagnetic tracking system (Liberty, Polhemus; 240 samples/s). A cursor representing this fingertip position was projected, together with the trial’s start and target positions, onto a back-projection screen in real-time (visual feedback delay ≤ 32 ms). The back-≤ projection screen was mounted horizontally above a first-surface mirror in such a manner that, when participants viewed the image in the mirror, the start position, target, and feedback cursor all appeared to be located in the same plane as the participant’s hand, even though the hand itself remained invisible below the mirror (Figure 7B).

**Figure 7.**
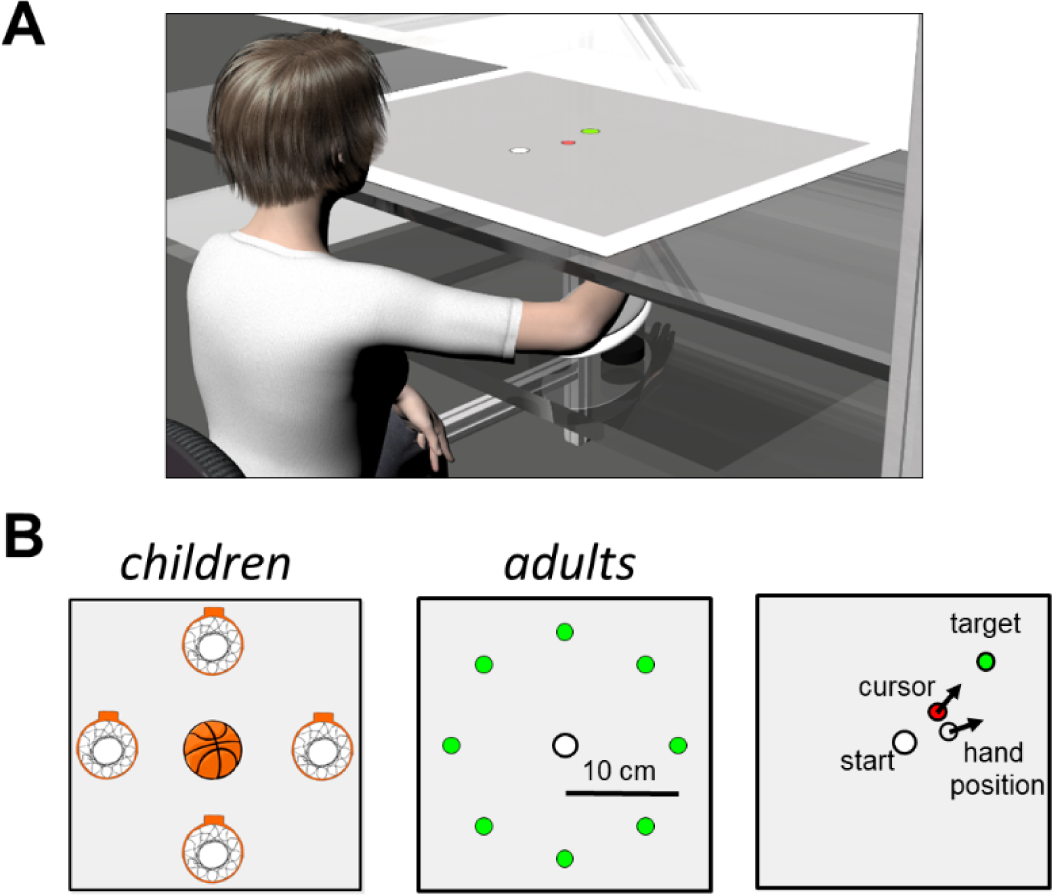
Experimental set-up for reach visuomotor adaptation. (A) Participant viewing the workspace displayed in a mirror that is part of a virtual display system. Reaching movements are performed with the right arm which is supported by an air sled on a glass surface. (B) The workspace displayed reach targets as basketball hoops for children (left) and as green circles for adults (middle). All possible targets are shown here, but targets appeared one at a time in blocked randomized order. Fast out-and-back reach movements were performed by moving a basketball (children) or red cursor (adults) to a target and back to the start position. During trials with perturbed visual feedback, the cursor’s position was rotated 30 degrees counter-clockwise around the center of the workspace (right).

#### Reach visuomotor adaptation task

Adult participants were instructed to make out-and-back reaching movements from a white center start position (radius 1 cm) to one of eight radially oriented green targets (radius 0.75 cm) located 10 cm from the start position at angles ranging from 0° to 315° in 45° increments. A custom MATLAB program presented all targets in random order within each block of 8 trials.

The start position was fixed in the participant’s midsagittal plane, with distance from the body adjusted such that the elbow was flexed approximately 90° when the fingertip rested on the start position. Continuous visual feedback of fingertip position was presented by a red cursor of the same size as the targets (radius 0.75 cm). Participants were asked to move the cursor out to the target and back to the start position without stopping at the target. They were told that it was important to move as fast as possible after initiating the movement, to not make any corrections during the movement, and that it was not necessary to initiate the movement quickly after the appearance of a new target.

Each session for adult participants consisted of 25 blocks of the 8 possible targets (200 trials) across three phases. During the *baseline* phase (initial 5 blocks, 40 trials), veridical feedback was presented (i.e., no perturbation). During the *full-exposure* phase (15 blocks, 120 trials), the position of the cursor representing fingertip position was rotated 30° counterclockwise (CCW) around the center of the work space (corresponding to the start position). During the wash-out or *after-effects* phase (5 blocks, 40 trials), veridical feedback was restored. Before data collection, each participant was given a set of familiarization trials with veridical feedback.

To test visuomotor adaptation in children, a shorter and child-friendly task was developed. Child participants were also instructed to make out-and-back reaching movements from a white center start position, but there were only four targets, located 10 cm from the start position at 0°, 90°, 180°, and 270°. The targets were represented by an image of a basketball hoop and the cursor was an image of a basketball (Figure 7B). The start position, cursor, and targets all had a radius of 2.5 cm. For each child, the start position location was determined in the same manner as described above for adults. Our custom MATLAB control program presented the targets in randomized order per block of 4 trials. We asked the children to move the ball to the hoop and back as fast as possible and without stopping at the hoop. They were encouraged throughout the task to keep moving fast and to not correct their movements within a trial even if the cursor missed the target.

Children completed 18 blocks of 4 targets (72 trials total). Like in the adults’ task, there were three phases: *baseline* (4 blocks, 16 trials), *full-exposure* (10 blocks, 40 trials), and *after-effects* (4 blocks, 16 trials). During the full-exposure phase, the position of the cursor representing fingertip position was rotated 30° CCW around the center of the workspace. During the baseline and after-effects phases, no perturbation was applied.

#### Data extraction and analyses

Custom MATLAB code smoothed the motion sensor data using a butterworth low-pass filter with cutoff frequency 10 Hz. These filtered position data were differentiated to obtain tangential velocity signals. Measurements of movement duration were made given that a between-group difference in this variable may allow the slower group to implement more within-trial corrections (as opposed to updating internal models relied upon during movement planning). Movement onset was defined as the time point where tangential velocity first exceeded 5 cm/s.

Movement offset was defined as the time point where tangential velocity dropped below 5 cm/s or, if that did not happen, where tangential velocity reached a local minimum.

For adult participants, initial reach angle was measured as the direction of a vector between start position and cursor location at the time point when peak tangential velocity was reached or 150 ms after movement onset, whichever occurred first (Tong & Flanagan, 2003). For each trial, this initial reach angle was then expressed relative to a vector from the start position to the specific target presented on that trial. Similar to the data processing steps in the speech auditory-motor adaptation task, all relative reach angles were normalized to baseline by subtracting, for each target direction separately, the median relative reach angle across baseline trials 3-5 for that same target. Lastly, the normalized reach angle for each block of 8 trials was obtained by averaging the normalized reach angles toward the 8 different targets in that block. For children, similar analysis procedures were used, but the cut-off for tangential peak velocity to be reached was 200 ms (rather than 150 ms) after movement onset. Thus, if tangential peak velocity had not been reached 200 ms after onset, we used the cursor location at that time point to calculate initial movement direction. Further, we used each target’s baseline trials 1-4 for normalizing the relative reach angles to baseline. Final data were obtained by averaging the normalized relative reach angles across the 4 different targets in each block of 4 trials. Trials were excluded if the tangential velocity did not exceed the threshold for movement onset (indicating no movement or an extremely slow movement) or if a movement clearly was not aimed at the target (such as when a child merely repeated a previous movement or turned to interact with the parent or experimenter). Overall, 0.3% of all trials for adult participants and 7.4% of all trials for child participants were excluded.

We conducted all statistical analyses once for final adaptation extent and once for initial rate of adaptation. To analyze adaptation extent, we used each participant’s last two blocks of the perturbation phase. For initial rate of adaptation, we fitted each participant’s data with an exponential function as in common in studies of reach adaptation (e.g., Flanagan et al., 1999; Smith et al., 2006). Specifically, we fitted the function 

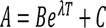

where *A* represents normalized relative reach angle data (averaged across 8 directions per block) during the perturbation phase and *T* represents the trial block. The unknown parameters *B*, λ, and *C* were determined by a nonlinear least squares algorithm computed using the *nls* function in R (R Core Team, 2018), and the λ values were used to quantify each individual’s initial rate of λ learning. However, the learning curves for most children were not exponential and, thus, not well fitted by the function. Hence, given that the function fitting method was only appropriate for adult participants, we also examined initial learning with an alternative method that determined how much adaptation had already occurred in the first two blocks of the perturbation phase.

For statistical analysis of the adult data, which involved only group as an independent variable, all between-group comparisons were completed with Welch’s *t*-tests in R (R Core Team, 2018). Cohen’s *d* effect sizes were calculated using the *cohen.d* function in the *effsize* package (Torchiano, 2018). For the children’s data, between-subjects independent variables included both group and age. We therefore used the *aov* function in R to conduct a two-way ANOVA to test both the stuttering vs. nonstuttering group difference as well as the differences across the age subgroups (3-4 years, 5-6 years, 7-9 years). Given the unbalanced design with a different number of children in each age subgroup, the type III sum of squares, *F*-values, and *p*-values of the ANOVA were calculated with the *Anova* function from the *Car* package (Fox & Weisberg, 2018). Effect sizes were calculated with the *MOTE* package (Buchanan et al., 2019).

## Results

### Movement duration

AWS showed descriptively longer movement durations than AWNS (*M* = 375 ms, *SD* = 119 ms, and *M* = 319 ms, *SD* = 41 ms, respectively), but this difference was not statistically significant, *t*(17.296) = -1.724, *p* = 0.103, *d* = 0.630. For children, there was also no statistically significant main effect for the between-group comparison of CWS and CWNS (*M* = 391 ms, *SD* = 58 ms, and *M* = 414 ms, *SD* = 98 ms, respectively), *F*(1, 48) = 0.943, *p* = 0.336, 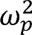 = -0.004.

There was a statistically significant main effect for age, *F*(2, 48) = 5.058, *p* = 0.010, 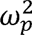 = 0.131, with movement duration getting shorter with increasing age (Age 3-4, *M* = 441 ms, *SD* = 78 ms; Age 5-6, *M* = 414 ms, *SD* = 90 ms; Age 7-9, *M* = 365 ms, *SD* = 56 ms), in the absence of a significant Group × Age interaction, *F*(2, 48) = 0.496, *p* = 0.612, 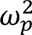 = -0.019.

### Visuomotor adaptation in AWS vs. AWNS

Adult participants in both the stuttering and nonstuttering groups exhibited adaptation in response to the visual perturbation. Figure 8A includes group average relative reach angle data throughout the baseline, perturbation, and after-effects phase. By the last two blocks of the perturbation phase, AWS and AWNS showed a similar final extent of adaptation that was not statistically different, *t*(21.569) = -0.598, *p* = 0.556, *d* = 0.218.

**Figure 8.**
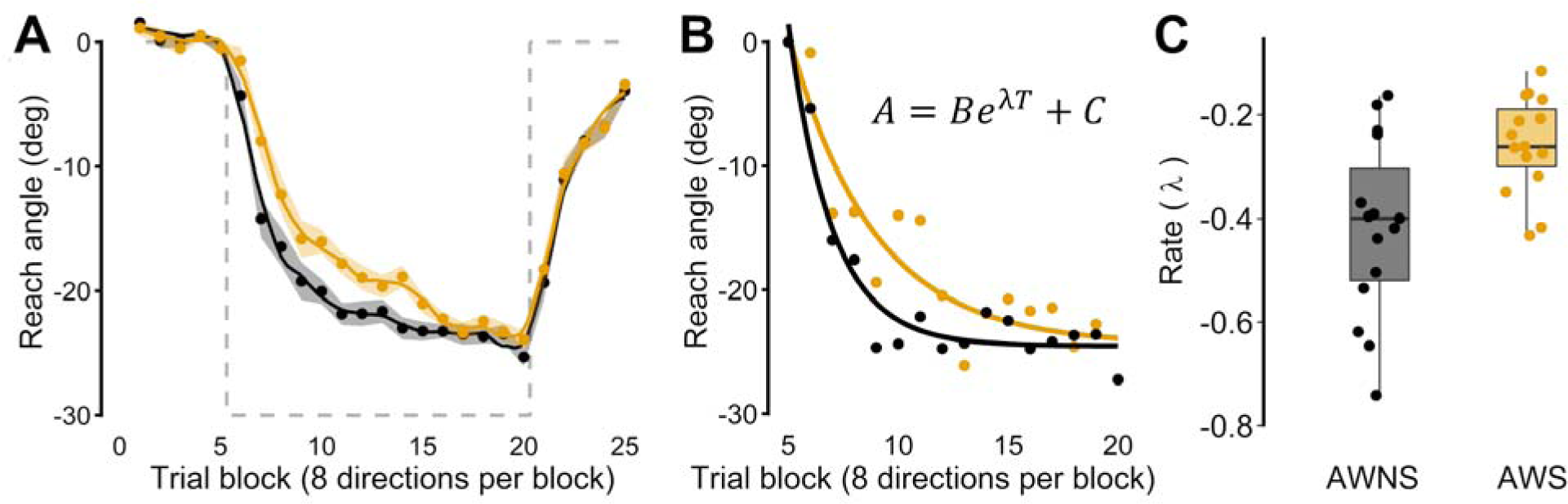
Visuomotor adaptation to a 30° CCW rotation for adults who do not stutter (black) and adults who stutter (orange). (A) Group averaged reach angles relative to baseline (averaged across 8 reach directions after normalization by direction). Shaded areas indicate standard errors of the mean. Gray dashed lines plot the visual perturbation with reversed sign (thus indicating hypothetical full compensation). (B) Two individual participant illustrations of exponential fitting of reach angle data from the perturbation phase. (C) Individual participant data for the exponential rate term (λ), overlaid on group box plots.

Given that we examined both learning rate (by fitting an exponential function) and early learning (by analyzing adaptation in the first two blocks of the perturbation phase), the associated between-group comparisons were treated as one family of two tests. Fitting an exponential function to each participant’s adaptation data (Figure 8B) revealed a statistically significant between-group difference in the initial rate of learning: the exponential rate term was significantly more negative (i.e., a faster drop in the curve and, thus, faster learning) for AWNS as compared with AWS, *t*(21.573) = -3.193, *p* = 0.009, *d* = 1.166 (Figure 8C). This result was confirmed by the second method of examining initial learning: the extent of adaptation reached in the first two blocks after introduction of the perturbation was significantly greater for AWNS than for AWS, *t*(25.430) = -3.033, *p* = 0.009, *d* = 1.108.

There was no statistically significant correlation between stuttering frequency and extent of visuomotor adaptation (*r*(13) = -0.008, *p* = 0.978). Stuttering frequency was also not significantly correlated with rate of adaptation determined by either the exponential rate term, *r*(13) = -0.375, *p* = 0.168, or the initial two perturbation blocks, *r*(13) = -0.085, *p* = 0.764.

### Visuomotor adaptation in CWS vs. CWNS

Both stuttering and nonstuttering children adapted to the perturbation (Figure 9), although the extent of adaptation was descriptively smaller (11-16*°* of the 30*°* perturbation) than that seen in adults (24-25*°* of the 30*°* perturbation). The ANOVA revealed that there was no statistically significant difference in final extent of adaptation between CWS and CWNS, *F*(1, 48) = 3.039, *p* = 0.088, 0.686, 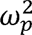 = 0.040. There was also no significant main effect of Age, *F*(2, 48) = 0.380, *p* =0.686, 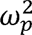 = -0.024. In addition, the Group × Age interaction was not significant, *F*(2, 48) = 0.444, *p* = 0.644, 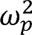 = -0.021. Similarly, the separate ANOVA for children’s initial extent of adaptation also indicated no statistically significant main effect of Group, *F*(1, 48) = 0.035, *p* = 0.852, 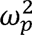 = -0.018, or Age, *F*(2, 48) = 0.034, *p* = 0.966, 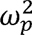 = -0.037, and no statistically significant interaction of Group and Age, *F*(2, 48) = 2.121, *p* = 0.131, 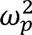 = 0.040.

**Figure 9.**
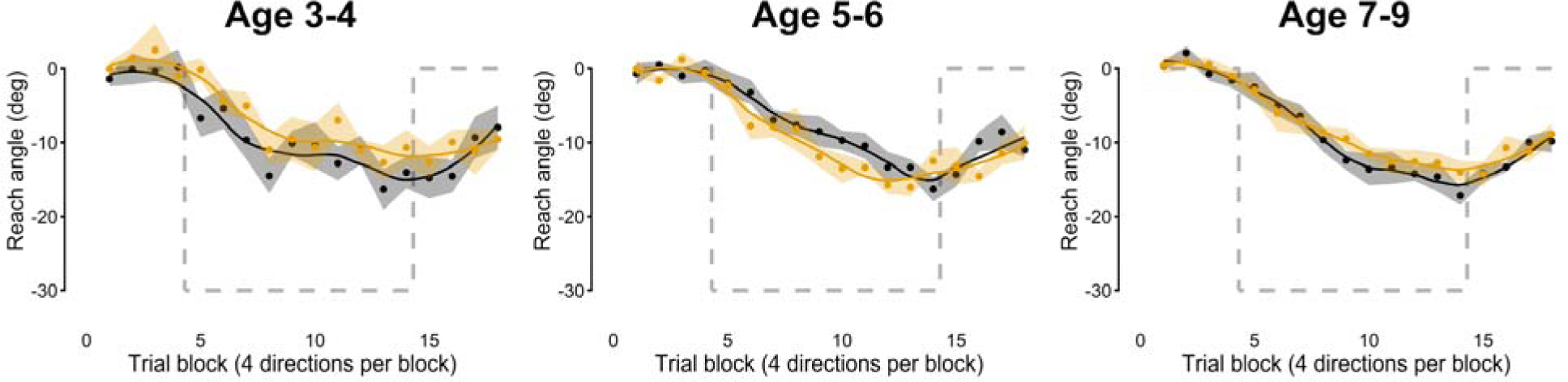
Visuomotor adaptation to a 30° CCW rotation for children who do not stutter (black) and children who stutter (orange). Data show group averaged reach angles relative to baseline (averaged across 4 reach directions after normalization by direction). Shaded areas indicate standard errors of the mean. Gray dashed lines plot the visual perturbation with reversed sign (thus indicating hypothetical full compensation).

Extent of adaptation in the final two blocks of the perturbation phase did not significantly correlate with age for either CWNS (*r*(25) = -0.049, *p* = 0.808) or CWS (*r*(25) = -0.091, *p* = 0.652). Similarly, initial adaptation in the first two blocks of the perturbation phase also did not significantly correlate with age for either CWNS (*r*(25) = 0.142, *p* = 0.480) or CWS (*r*(25) = -0.109, *p* = 0.588). Lastly, for none of the three age groups of CWS was there a statistically significant correlation between the final extent of visuomotor adaptation and stuttering frequency from the clinical assessment (Age 3-4, *r*(7) = - 0.040, *p* = 0.919; Age 5-6, *r*(6) = 0.028, *p* = 0.948; Age 7-9, *r*(8) = 0.426, *p* = 0.220), or between initial visuomotor adaptation and stuttering frequency from the clinical assessment (Age 3-4, *r*(7) = -0.364, *p* = 0.335; Age 5-6, *r*(6) = 0.162, *p* = 0.702; Age 7-9, *r*(8) = 0.465, *p* = 0.176).

## Discussion

We investigated nonspeech sensorimotor learning in adults and children who stutter as compared with age-matched individuals who do not stutter. Specifically, participants were tested in a visuomotor rotation paradigm that involved making fast out-and-back reaching movements while visual feedback was manipulated during the perturbation phase of the experiment. The perturbation involved a 30° CCW rotation around the center of the workspace (which was also the start position for all movements) of the location of a cursor representing fingertip position.

The participant’s hand and arm remained hidden below a screen throughout the experiment. Analyses focused on participants’ adaptation to this perturbation (i.e., adjustments in reach direction in the opposite direction of the perturbation) in terms of both overall adaptation extent and initial rate of adaptation.

For adults, findings show that AWS did reach the same final level of adaptation as AWNS after more than 80 exposure trials, but that their initial rate of learning was statistically significantly slower. Thus, stuttering adults’ limitations in limb visuomotor learning are more subtle than those observed in the speech auditory-motor learning task in Experiment 1 where their overall extent of learning was also limited. Nevertheless, this novel finding of a lower initial rate of learning in adjusting the planning of upper limb reach movements suggests that stuttering individuals’ sensorimotor integration difficulties are not restricted to either the orofacial effector system or speech-specific sensorimotor control processes.

Unlike the situation for speech auditory-motor adaptation, the reach visuomotor adaptation task revealed a between-group difference only for adults and not for children. Indeed, CWS did not differ from CWNS in either the final extent of learning or the initial rate of learning in any of the three age groups (3-4, 5-6, 7-9 years of age). This result could be interpreted as suggesting that nonspeech visuomotor learning deficits develop only after the age of 9 years, which is typically several years after the onset of stuttering. Given that it is not obvious how stuttering itself, personal experiences related to stuttering, or techniques applied in stuttering treatment could contribute to the development of upper limb motor learning difficulties, the mechanisms behind such late-developing nonspeech effects would also remain entirely unknown.

Alternatively, it is possible that the different outcome for speech auditory-motor learning (differences with control participants were observed in both AWS and CWS) as opposed to reach visuomotor learning (differences with control participants were observed in AWS but not CWS) stems from inherent differences in the typical developmental trajectories for these two tasks. In our speech auditory-motor adaptation task, even the youngest group of 3-4-year-old nonstuttering children already showed an extent of adaptation that was similar to that of the nonstuttering adult participants (although the size of the perturbation and number of exposure trials differed between the child and adult tasks, under those different experimental circumstances CWNS’ extent of adaptation nevertheless reached the same magnitude as that of AWNS). Thus, speech auditory-motor learning in typically developing children appears to follow a fast developmental trajectory, and an advanced learning ability in this domain may be necessary from an early age to correctly update the internal models that take account of how maturational changes in the biomechanical and neural systems for speech continually affect the transformations from motor commands to acoustic speech output (see Callan et al., 2000; Kent, 1976; Kent, 1997; Vorperian et al., 1999; Vorperian et al., 2005; Vorperian & Kent, 2007; Vorperian et al., 2009). In contrast, for our child and adult reach visuomotor adaptation tasks (again acknowledging a number of methodological differences such as number of exposure trials and number of different targets), even the oldest group of 7-9-year-old nonstuttering children reached only approximately 65% of the extent of adaptation observed for the nonstuttering adults. Although resulting here from non-identical child and adult tasks, this descriptive difference is consistent with prior work demonstrating through direct age-group comparisons that several aspects of upper and lower limb sensorimotor learning continue to develop throughout the school-age years (Contreras-Vidal et al., 2005; Ferrel et al., 2001; Kagerer & Clark, 2014; Rossi et al., 2019). Moreover, Kagerer and Clark (2015) specifically reported that the learning of visuomotor maps may continue to develop gradually throughout childhood whereas the learning of (nonspeech) auditory-motor maps may reach adult-like performance levels around the age of 5-6 years. Taking together our new findings and the results from the cited studies, upper limb visuomotor learning shows a more protracted developmental trajectory, and even typical children in the age range tested here (3-9 years of age) are still limited in their learning abilities within this domain. As a result, subtle nonspeech sensorimotor learning differences between stuttering and nonstuttering individuals may not become detectable until after the latter group reaches adult levels of performance.

In sum, our specific visuomotor adaptation task identified a difference in sensorimotor learning for fast reaching movements in stuttering vs. nonstuttering adults but not in stuttering vs. nonstuttering children. The between-group difference for adult participants was found in the initial rate of learning, with AWS learning more slowly, and, thus, requiring more trials with perturbed visual feedback before achieving the same level of adaptation as their nonstuttering peers. By the end of the task, however, both groups showed equivalent levels of learning, and their adjusted reach directions showed similar degrees of compensation for the perturbation. In children, neither initial nor overall learning differentiated between CWS and age-matched nonstuttering children for three different age groups. Interestingly, under their respective experimental conditions, even CWNS showed limited visuomotor learning in comparison with AWNS (whereas for speech auditory-motor adaptation in Experiment 1, CWNS showed similar learning as AWNS). Considered in the context of the above referenced literature (Contreras-Vidal et al., 2005; Ferrel et al. 2001; Kagerer & Clark, 2014, 2015; Rossi et al., 2019), we tentatively interpret these nonspeech findings as suggesting that (a) the typical developmental trajectory for upper limb visuomotor learning is relatively slow in comparison with speech production, (b) this slower trajectory for typical development may indicate that fully mature learning abilities are not required for upper limb sensorimotor control during childhood, and (c) therefore, the CNS of children who stutter is able to appropriately meet these nonspeech sensorimotor learning demands.

## General discussion

The aim of this work was to compare CWS and CWNS as well as AWS and AWNS with regard to both speech auditory-motor learning and reach visuomotor learning. Age-, sex-, and handedness-matched stuttering and nonstuttering participants were tested in a speech adaptation task with a real-time perturbation of the formant frequencies in the auditory feedback or a reach adaptation task with a real-time perturbation of the cursor location in the visual feedback.

Acoustic recordings of speech were used to determine to what extent participants learned to adjust their formant frequencies in the speech task, and motion tracking of the fingertip was used to determine the rate and extent of adjustments in movement direction in the reach task.

Experiment 1 yielded several interesting findings. AWS showed limitations in speech auditory-motor learning in response to both upward and downward formant perturbations, and this was true for both suddenly and gradually introduced perturbations. Interestingly, the observed difference between stuttering and nonstuttering adults was larger for perturbations that were introduced all-at-once in-between two successive trials (the sudden condition) as compared with perturbations that were incrementally ramped up over many trials (the gradual condition). Whether or not this more severe impact on learning in the sudden condition than in the gradual condition can inform on the relative involvement of cortico-cortical, basal ganglia, and cerebellar learning mechanisms in stuttering remains open for debate. Although it is well documented that stuttering is associated with structural and functional differences in sensorimotor fronto-parieto-temporal pathways, basal ganglia, and cerebellum (Alm, 2004; Anderson et al., 1999; Brown et al., 2005; Chang & Zhu, 2013; Civier et al., 2013; De Nil et al., 2001; Fox, P. T. et al., 1996; Giraud et al., 2008; Koller, 1983; Lu et al., 2010; Maguire et al., 2012; Shahed & Jankovic, 2001; Wu et al., 1995; Wu et al., 1997), the differential involvement of these neural substrates in various sensorimotor adaptation paradigms has remained elusive (Contreras-Vidal & Buch, 2003; e.g., Criscimagna-Hemminger et al., 2010; Doya, 2000; Doyon et al., 2003; Gutierrez-Garralda et al., 2013; Maschke et al., 2004; Rabe et al., 2009; Robertson & Miall, 1999; Smith & Shadmehr, 2005; Venkatakrishnan et al., 2011; Weiner et al., 1983; Werner et al., 2009; Werner et al., 2010).

Probably the most important finding from Experiment 1 is that a limitation in auditory-motor learning is also present in young CWS. Statistically, neither of the two age groups of CWS showed any learning in either the sudden or the gradual condition. In the same conditions, age-matched CWNS showed a large extent of learning that was similar to that seen in typical adults. As a related novel finding, CWS’ extent of speech auditory-motor learning was positively correlated with age whereas CWNS did not show such a relationship. In other words, these new correlational findings suggest that the capacity for auditory-motor learning was still developing between the ages of 3 and 9 years in CWS, but had already reached an adult-like level by the age of 3 years in CWNS. More generally, our overall results (considering the statistical significance and effect sizes of between-group differences together with the correlational results) provide compelling evidence in favor of the view that speech auditory-motor learning impairments are already present, and in fact more severe, in the early childhood years closest to the onset of stuttering. Hence, these limitations certainly have the potential to be a reflection of mechanisms directly involved in the proximal cause of stuttering rather than merely being a by-product of years of experience with stuttering.

Experiment 2 revealed that sensorimotor learning limitations in AWS are also present, albeit in more subtle form, in an upper limb reaching task. In the case of this visuomotor learning task, there was only a difference between AWS and AWNS in the initial rate of learning and not in the total extent of learning achieved by the end of the task. In contrast, CWS did not differ from CWNS in either initial or final amount of visuomotor adaptation. This discrepancy in the nonspeech results for children vs. adults should be considered in light of the fact that even CWNS showed much less adaptation than AWNS. It is therefore possible that subtle visuomotor learning difficulties may not be detectable with the present nonspeech task until nonstuttering individuals achieve mature learning levels.

Taken together, the findings from Experiment 1 and Experiment 2 suggest that CWS’ developmental trajectory for sensorimotor learning may be sufficiently fast to appropriately adjust motor commands for upper limb movements but not those for orofacial speech movements. That is, the daily-life demands on upper limb visuomotor adaptation (i.e., adjusting internal models to take account of neural and limb biomechanical changes that affect motor-to-sensory transformations) may be *relatively* low given that our data (as well as those of several other laboratories cited above) show that even 7-9-year-old CWNS still have limited learning abilities in this domain, yet they typically do not show clinically relevant upper limb movement problems. On the other hand, the demands on speech auditory-motor adaptation (i.e., adjusting internal models to take account of neural and orofacial biomechanical changes that affect motor-to-sensory but also vocal tract-to-acoustics transformations) may be more stringent given that our data show that typical development in this domain already achieves adult-like learning abilities as early as 3-6 years of age. As we found the development of speech auditory-motor learning abilities to be slower in children who stutter, their delay in this domain may be detrimental to fluent speech production. Thus, overall, findings from the present speech adaptation task are consistent with our previous proposal that children who stutter are limited in their ability to update the control policy (internal inverse model) and/or internal forward model that are critical for speech sensorimotor control (Max & Daliri, 2019; Max, 2004; Max et al., 2004).

An important challenge for future studies will be to directly test the related proposal that some sensory prediction errors resulting from movement planning with poorly updated internal models (Flanagan et al., 2003; Haith & Krakauer, 2013; Wolpert & Miall, 1996; Wolpert & Flanagan, 2001) can indeed trigger maladaptive system corrections that involve repeated movements or fixed postures. Additional pivotal insights may be gained from efforts investigating which specific sensory, motor, or learning mechanisms impede sensorimotor learning—and in particular auditory-motor learning—in children and adults who stutter. In our own lab, we have recently been able to demonstrate that adults who stutter show deficits in at least some aspects of the sensory prediction mechanism that drives internal model learning (Daliri & Max, 2015a; Daliri & Max, 2015b; Daliri & Max, 2018; Max & Daliri, 2019). The acuity of sensory processing itself during and after movement execution also warrants further investigation. Despite multiple reports of reduced oral somatosensory acuity in individuals who stutter (Archibald & De Nil, 1999; De Nil & Abbs, 1991; Howell et al., 1995; Loucks & De Nil, 2006a; Loucks & De Nil, 2006b; although see Daliri et al., 2013), auditory acuity for the detection of unpredictable, within-trial formant frequency perturbations was reported to not differ between stuttering and nonstuttering participants (Cai et al., 2012). However, the isolated vowel productions tested in the latter study were substantially longer (∼300 ms in duration) than those occurring in connected speech, and this additional time may have been beneficial for stuttering speakers to detect the formant changes. Lastly, for upper limb movements, it has become increasingly clear that the overall process of sensorimotor learning comprises dissociable implicit and explicit learning components, with the former being driven by prediction error and the latter by performance error (Mazzoni & Krakauer, 2006; McDougle et al., 2015; Schween & Hegele, 2017; Taylor et al., 2014). To date, several lines of evidence suggest that speech auditory-motor learning is predominantly implicit in nature (see Shiller et al., 2020).

Thus, our combined speech auditory-motor and reach visuomotor learning results are consistent with the idea that individuals who stutter have difficulty specifically with the implicit component of sensorimotor learning. This hypothesis can be tested directly in future speech experiments that adopt novel methodological approaches from the limb motor control literature (Mazzoni & Krakauer, 2006; McDougle et al., 2015; Schween & Hegele, 2017; Taylor et al., 2014).

## Acknowledgements

Roman Prokopenko and Caitlin Baldwin made major contributions to data collection and analysis. Derek Maffett, Churan Li, Patrick McCleary, Elizabeth Rylance, and Prince Z. Wang also assisted with data collection or analysis. Allison Stewart, Annie Zhu, and Chrystal Chau conducted speech and language evaluations and stuttering severity assessments. This research was supported by grants from the National Institute on Deafness and Other Communication Disorders (R01 DC007603, R01 DC014510, and R01 DC17444, L. Max PI) and the Canadian Institutes of Health Research (MOP-137001, L. Max Co-I). The content is solely the responsibility of the authors and does not necessarily represent the official views of the National Institute on Deafness and Other Communication Disorders, the National Institutes of Health, or the Canadian Institutes of Health Research.

## Footnotes

1 Due to a communication malfunction between the control computer and vocal effects processor, one adult and one child from the stuttering groups experienced a delay of 35 ms instead of 10 ms. As the amount of learning observed for these two participants was typical for that seen in their respective groups, we included the obtained data sets and also applied an identical 35 ms delay during the recording sessions of the two matched nonstuttering participants.

2 Adult participants’ auditory-motor adaptation task included separate conditions with upward and downward perturbation directions, so we normalized the data by dividing by 250 for down-shift conditions and by -250 for up-shift conditions. This caused adaptive changes (i.e., moving in the opposite direction of the perturbation) to be represented as a positive number in the range from 0 to 1, with 1 corresponding to full compensation. Following the perturbation (i.e., moving in the same direction of the perturbation) resulted in a negative number.

## References

Alm, P. A. (2004). Stuttering and the Basal Ganglia Circuits: A Critical Review of Possible Relations. Journal of Communication Disorders, 37(4), 325–369. https://doi.org/10.1016/j.jcomdis.2004.03.001

Anderson, J. M., Hughes, J. D., Rothi, L. J., Crucian, G. P., & Heilman, K. M. (1999). Developmental Stuttering and Parkinson’s Disease: The Effects of Levodopa Treatment. Journal of Neurology, Neurosurgery, and Psychiatry, 66(6), 776–778. https://doi.org/10.1136/jnnp.66.6.776

Archibald, L., & De Nil, L.,F. (1999). The Relationship between Stuttering Severity and Kinesthetic Acuity for Jaw Movements in Adults Who Stutter. Journal of Fluency Disorders, 24(1), 25–42. https://doi.org/10.1016/S0094-730X(98)00023-0

Beal, D. S., Gracco, V. L., Brettschneider, J., Kroll, R. M., & De Nil, L. F. (2013). A Voxel-Based Morphometry (VBM) Analysis of Regional Grey and White Matter Volume Abnormalities within the Speech Production Network of Children Who Stutter. Cortex; a Journal Devoted to the Study of the Nervous System and Behavior, 49(8), 2151–2161. https://doi.org/10.1016/j.cortex.2012.08.013

Benito-Aragón, C., Gonzalez-Sarmiento, R., Liddell, T., Diez, I., d’Oleire Uquillas, F., Ortiz-Terán, L., Bueichekú, E., Chow, H. M., Chang, S., & Sepulcre, J. (2020). Neurofilament-Lysosomal Genetic Intersections in the Cortical Network of Stuttering. Progress in Neurobiology, 184, 101718. https://doi.org/10.1016/j.pneurobio.2019.101718

Bishop, J. H., Williams, H. G., & Cooper, W. A. (1991). Age and Task Complexity Variables in Motor Performance of Children with Articulation-Disordered, Stuttering, and Normal Speech. Journal of Fluency Disorders, 16(4), 219–228. https://doi.org/10.1016/0094-730X(91)90004-V

Boersma, P., & Weenink, D. (2008). Praat: doing phonetics by computer [computer software]

Brown, S., Ingham, R. J., Ingham, J. C., Laird, A. R., & Fox, P. T. (2005). Stuttered and Fluent Speech Production: An ALE Meta-Analysis of Functional Neuroimaging Studies. Human Brain Mapping, 25(1), 105–117. https://doi.org/10.1002/hbm.20140

Buchanan, E. M., Gillenwaters, A., Scofield, J. E., & Valentine, K. D. (2019). MOTE: Measure of the Effect: Package to Assist in Effect Size Calculations and their Confidence Intervals. http://github.com/doomlab/MOTE.

Cai, S., Beal, D. S., Ghosh, S. S., Tiede, M. K., Guenther, F. H., & Perkell, J. S. (2012). Weak Responses to Auditory Feedback Perturbation during Articulation in Persons Who Stutter: Evidence for Abnormal Auditory-Motor Transformation. PloS One, 7(7), e41830. https://doi.org/10.1371/journal.pone.0041830

Cai, S., Tourville, J. A., Beal, D. S., Perkell, J. S., Guenther, F. H., & Ghosh, S. S. (2014). Diffusion Imaging of Cerebral White Matter in Persons Who Stutter: Evidence for Network-Level Anomalies. Frontiers in Human Neuroscience, 8 https://doi.org/10.3389/fnhum.2014.00054

Callan, D. E., Kent, R. D., Guenther, F. H., & Vorperian, H. K. (2000). An Auditory-Feedback Based Neural Network Model of Speech Production that is Robust to Developmental Changes in the Size and Shape of the Articulatory System. Journal of Speech, Language, and Hearing Research, 43, 721–736. https://doi.org/10.1044/jslhr.4303.721

Chang, S., Horwitz, B., Ostuni, J., Reynolds, R., & Ludlow, C. L. (2011). Evidence of Left Inferior Frontal-Premotor Structural and Functional Connectivity Deficits in Adults Who Stutter. Cerebral Cortex (New York, N.Y.: 1991), 21(11), 2507-2518. https://doi.org/10.1093/cercor/bhr028

Chang, S., & Zhu, D. C. (2013). Neural Network Connectivity Differences in Children Who Stutter. Brain : A Journal of Neurology, 136(Pt 12), 3709–3726. https://doi.org/10.1093/brain/awt275

Choo, A. L., Chang, S., Zengin-Bolatkale, H., Ambrose, N. G., & Loucks, T. M. (2012). Corpus Callosum Morphology in Children Who Stutter. Journal of Communication Disorders, 45(4), 279–289. https://doi.org/10.1016/j.jcomdis.2012.03.004

Civier, O., Bullock, D., Max, L., & Guenther, F. H. (2013). Computational Modeling of Stuttering Caused by Impairments in a Basal Ganglia Thalamo-Cortical Circuit Involved in Syllable Selection and Initiation. Brain and Language, 126(3), 263–278. https://doi.org/10.1016/j.bandl.2013.05.016

Contreras-Vidal, J. L., Bo, J., Boudreau, J. P., & Clark, J. E. (2005). Development of Visuomotor Representations for Hand Movement in Young Children. Experimental Brain Research, 162(2), 155–164. https://doi.org/10.1007/s00221-004-2123-7

Contreras-Vidal, J. L., & Buch, E. R. (2003). Effects of Parkinson’s Disease on Visuomotor Adaptation. Experimental Brain Research, 150(1), 25–32. https://doi.org/10.1007/s00221-003-1403-y

Coren, S. (1993). The Lateral Preference Inventory for Measurement of Handedness, Footedness, Eyedness, and Earedness: Norms for Young Adults. Bulletin of the Psychonomic Society, 31(1), 1–3. https://doi.org/10.3758/BF03334122

Cornelisse, L. E., Gagné, J., & Seewald, R. C. (1991). Ear Level Recordings of the Long-Term Average Spectrum of Speech*. Ear and Hearing, 12(1), 47.

Criscimagna-Hemminger, S. E., Bastian, A. J., & Shadmehr, R. (2010). Size of Error Affects Cerebellar Contributions to Motor Learning. Journal of Neurophysiology, https://doi.org/10.1152/jn.00822.2009

Daliri, A., & Max, L. (2015a). Electrophysiological Evidence for a General Auditory Prediction Deficit in Adults Who Stutter. Brain and Language, 150, 37–44. https://doi.org/10.1016/j.bandl.2015.08.008

Daliri, A., & Max, L. (2015b). Modulation of Auditory Processing during Speech Movement Planning is Limited in Adults Who Stutter. Brain and Language, 143, 59–68. https://doi.org/10.1016/j.bandl.2015.03.002

Daliri, A., & Max, L. (2018). Stuttering Adults’ Lack of Pre-Speech Auditory Modulation Normalizes when Speaking with Delayed Auditory Feedback. Cortex; a Journal Devoted to the Study of the Nervous System and Behavior, 99, 55–68. https://doi.org/10.1016/j.cortex.2017.10.019

Daliri, A., Prokopenko, R. A., Flanagan, J. R., & Max, L. (2014). Control and Prediction Components of Movement Planning in Stuttering Versus Nonstuttering Adults. Journal of Speech, Language, and Hearing Research: JSLHR, 57(6), 2131–2141. https://doi.org/10.1044/2014_JSLHR-S-13-0333

Daliri, A., Prokopenko, R. A., & Max, L. (2013). Afferent and Efferent Aspects of Mandibular Sensorimotor Control in Adults Who Stutter. Journal of Speech, Language, and Hearing Research: JSLHR, 56(6), 1774–1788. https://doi.org/10.1044/1092-4388(2013/12-0134)

Daliri, A., Wieland, E. A., Cai, S., Guenther, F. H., & Chang, S. (2018). Auditory-Motor Adaptation is Reduced in Adults Who Stutter but Not in Children Who Stutter. Developmental Science, 21(2)https://doi.org/10.1111/desc.12521

De Nil, L. F., & Abbs, J. H. (1991). Kinaesthetic Acuity of Stutterers and Non-Stutterers for Oral and Non-Oral Movements. Brain: A Journal of Neurology, 114 ( *Pt 5*), 2145–2158. http://www.ncbi.nlm.nih.gov/pubmed/1933239

De Nil, L. F., Kroll, R. M., & Houle, S. (2001). Functional Neuroimaging of Cerebellar Activation during Single Word Reading and Verb Generation in Stuttering and Nonstuttering Adults. Neuroscience Letters, 302(2-3), 77–80. https://doi.org/10.1016/s0304-3940(01)01671-8

Desmurget, M., & Grafton, S. (2000). Forward Modeling Allows Feedback Control for Fast Reaching Movements. Trends in Cognitive Sciences, 4(11), 423–431. https://doi.org/10.1016/s1364-6613(00)01537-0

Doya, K. (2000). Complementary Roles of Basal Ganglia and Cerebellum in Learning and Motor Control. Current Opinion in Neurobiology, 10(6), 732–739. https://doi.org/10.1016/s0959-4388(00)00153-7

Doyon, J., Penhune, V., & Ungerleider, L. G. (2003). Distinct Contribution of the Cortico-Striatal and Cortico-Cerebellar Systems to Motor Skill Learning. Neuropsychologia, 41(3), 252–262. https://doi.org/10.1016/s0028-3932(02)00158-6

Dunn, L. M., & Dunn, D. M. (2007). PPVT-4: Peabody picture vocabulary test. Pearson Assessments.

Feng, Y., Gracco, V., & Max, L. (2011). Integration of Auditory and Somatosensory Error Signals in the Neural Control of Speech Movements. Journal of Neurophysiology, 106(2), 667–679. https://doi.org/10.1152/jn.00638.2010

Ferrel, C., Bard, C., & Fleury, M. (2001). Coordination in Childhood: Modifications of Visuomotor Representations in 6- to 11-Year-Old Children. Experimental Brain Research, 138(3), 313–321. https://doi.org/10.1007/s002210100697

Flanagan, J. R., Nakano, E., Imamizu, H., Osu, R., Yoshioka, T., & Kawato, M. (1999). Composition and Decomposition of Internal Models in Motor Learning Under Altered Kinematic and Dynamic Environments. The Journal of Neuroscience: The Official Journal of the Society for Neuroscience, 19(20), RC34. https://doi.org/10.1523/JNEUROSCI.19-20-j0005.1999

Flanagan, J. R., Vetter, P., Johansson, R. S., & Wolpert, D. M. (2003). Prediction Precedes Control in Motor Learning. Current Biology: CB, 13(2), 146–150. https://doi.org/10.1016/s0960-9822(03)00007-1

Forster, D. C., & Webster, W. G. (2001). Speech-Motor Control and Interhemispheric Relations in Recovered and Persistent Stuttering. Developmental Neuropsychology, 19(2), 125–145. https://doi.org/10.1207/S15326942DN1902_1

Fox, J., & Weisberg, S. (2018). An R Companion to Applied Regression (Third ed.). SAGE.

Fox, P. T., Ingham, R. J., Ingham, J. C., Hirsch, T. B., Downs, J. H., Martin, C., Jerabek, P., Glass, T., & Lancaster, J. L. (1996). A PET Study of the Neural Systems of Stuttering. Nature, 382(6587), 158–161. https://doi.org/10.1038/382158a0

Frigerio-Domingues, C., & Drayna, D. (2017). Genetic Contributions to Stuttering: The Current Evidence. Molecular Genetics & Genomic Medicine, 5(2), 95–102. https://doi.org/10.1002/mgg3.276

Frigerio-Domingues, C., Gkalitsiou, Z., Zezinka, A., Sainz, E., Gutierrez, J., Byrd, C., Webster, R., & Drayna, D. (2019). Genetic Factors and Therapy Outcomes in Persistent Developmental Stuttering. Journal of Communication Disorders, 80, 11–17. https://doi.org/10.1016/j.jcomdis.2019.03.007

Giraud, A. L., Neumann, K., Bachoud-Levi, A. C., von Gudenberg, A. W., Euler, H. A., Lanfermann, H., & Preibisch, C. (2008). Severity of Dysfluency Correlates with Basal Ganglia Activity in Persistent Developmental Stuttering. Brain and Language, 104(2), 190–199. https://doi.org/10.1016/j.bandl.2007.04.005

Goldman, R., & Fristoe, M. (2000). Goldman Fristoe Test of Articulation-2 (GFTA-2). American Guidance Service. Inc.

Goldman, R., & Fristoe, M. (2015). GFTA-3: Goldman Fristoe 3 Test of Articulation. PsychCorp.

Gutierrez-Garralda, J. M., Moreno-Briseño, P., Boll, M., Morgado-Valle, C., Campos-Romo, A., Diaz, R., & Fernandez-Ruiz, J. (2013). The Effect of Parkinson’s Disease and Huntington’s Disease on Human Visuomotor Learning. The European Journal of Neuroscience, 38(6), 2933–2940. https://doi.org/10.1111/ejn.12288

Haith, A. M., & Krakauer, J. W. (2013). Model-Based and Model-Free Mechanisms of Human Motor Learning. Advances in Experimental Medicine and Biology, 782, 1–21. https://doi.org/10.1007/978-1-4614-5465-6_1

Holm, S. (1979). A Simple Sequentially Rejective Multiple Test Procedure. Scandinavian Journal of Statistics, 6(2), 65–70. http://www.jstor.org/stable/4615733

Howell, P., Sackin, S., & Rustin, L. (1995). Comparison of Speech Motor Development in Stutterers and Fluent Speakers between 7 and 12 Years Old. Journal of Fluency Disorders, 20(3), 243–255. https://doi.org/10.1016/0094-730X(94)00011-H

Hresko, W. P., Reid, D. K., & Hammill, D. D. (1999). TELD-3: Test of early language development. Pro-ed.

Ingham, R. J., Fox, P. T., Ingham, J. C., Xiong, J., Zamarripa, F., Hardies, L. J., & Lancaster, J. L. (2004). Brain Correlates of Stuttering and Syllable Production: Gender Comparison and Replication. Journal of Speech, Language, and Hearing Research: JSLHR, 47(2), 321–341. https://doi.org/10.1044/1092-4388(2004/026)

Jones, R. D., White, A. J., Lawson, K. H. C., & Anderson, T. J. (2002). Visuoperceptual and Visuomotor Deficits in Developmental Stutterers: An Exploratory Study. Human Movement Science, 21(5), 603–619. https://doi.org/10.1016/S0167-9457(02)00165-3

Kagerer, F. A., & Clark, J. E. (2014). Development of Interactions between Sensorimotor Representations in School-Aged Children. Human Movement Science, 34, 164–177. https://doi.org/10.1016/j.humov.2014.02.001

Kagerer, F. A., & Clark, J. E. (2015). Development of Kinesthetic-Motor and Auditory-Motor Representations in School-Aged Children. Experimental Brain Research, 233(7), 2181–2194. https://doi.org/10.1007/s00221-015-4288-7

Kang, C., Riazuddin, S., Mundorff, J., Krasnewich, D., Friedman, P., Mullikin, J. C., & Drayna, D. (2010). Mutations in the Lysosomal Enzyme-Targeting Pathway and Persistent Stuttering. The New England Journal of Medicine, 362(8), 677–685. https://doi.org/10.1056/NEJMoa0902630

Kawato, M. (1999). Internal Models for Motor Control and Trajectory Planning. Current Opinion in Neurobiology, 9(6), 718–727. https://doi.org/10.1016/s0959-4388(99)00028-8

Kent, R. D. (1976). Anatomical and Neuromuscular Maturation of the Speech Mechanism: Evidence from Acoustic Studies. Journal of Speech and Hearing Research, 19(3), 421–447.

Kent, R. D. (1997). The speech sciences. Singular.

Kim, K. S., Wang, H., & Max, L. (2020). It’s about Time: Minimizing Hardware and Software Latencies in Speech Research with Real-Time Auditory Feedback. Journal of Speech, Language, and Hearing Research, 63(8), 2522–2534. https://doi.org/10.1044/2020_JSLHR-19-00419

Koller, W. C. (1983). Dysfluency (Stuttering) in Extrapyramidal Disease. Archives of Neurology, 40(3), 175–177. https://doi.org/10.1001/archneur.1983.04050030069014

Krakauer, J. W., & Mazzoni, P. (2011). Human Sensorimotor Learning: Adaptation, Skill, and Beyond. Current Opinion in Neurobiology, 21(4), 636–644. https://doi.org/10.1016/j.conb.2011.06.012

Loucks, T. M., & De Nil, L. F. (2006a). Anomalous Sensorimotor Integration in Adults Who Stutter: A Tendon Vibration Study. Neuroscience Letters, 402(1-2), 195–200. https://doi.org/10.1016/j.neulet.2006.04.002

Loucks, T. M., & De Nil, L. F. (2006b). Oral Kinesthetic Deficit in Adults Who Stutter: A Target-Accuracy Study. Journal of Motor Behavior, 38(3), 238–246. https://doi.org/10.3200/JMBR.38.3.238-247

Lu, C., Chen, C., Ning, N., Ding, G., Guo, T., Peng, D., Yang, Y., Li, K., & Lin, C. (2010). The Neural Substrates for Atypical Planning and Execution of Word Production in Stuttering. Experimental Neurology, 221(1), 146–156. https://doi.org/10.1016/j.expneurol.2009.10.016

Maguire, G. A., Yeh, C. Y., & Ito, B. S. (2012). Overview of the Diagnosis and Treatment of Stuttering. Journal of Experimental & Clinical Medicine, 4(2), 92–97. https://doi.org/10.1016/j.jecm.2012.02.001

Maschke, M., Gomez, C. M., Ebner, T. J., & Konczak, J. (2004). Hereditary Cerebellar Ataxia Progressively Impairs Force Adaptation during Goal-Directed Arm Movements. Journal of Neurophysiology, 91(1), 230–238. https://doi.org/10.1152/jn.00557.2003

Max, L. (2004). Stuttering and internal models for sensorimotor control: A theoretical perspective to generate testable hypotheses. Speech Motor Control in Normal and Disordered Speech (pp. 357–388). Oxford University Press.

Max, L., & Daliri, A. (2019). Limited Pre-Speech Auditory Modulation in Individuals Who Stutter: Data and Hypotheses. Journal of Speech, Language, and Hearing Research: JSLHR, 62(8S), 3071–3084. https://doi.org/10.1044/2019_JSLHR-S-CSMC7-18-0358

Max, L., Guenther, F. H., Gracco, V. L., Ghosh, S. S., & Wallace, M. E. (2004). Unstable Or Insufficiently Activated Internal Models and Feedback-Biased Motor Control as Sources of Dysfluency: A Theoretical Model of Stuttering. Contemporary Issues in Communication Sciences and Disorders, 31, 105–122. https://doi.org/10.1044/cicsd_31_S_105

Max, L., & Maffett, D. G. (2015). Feedback Delays Eliminate Auditory-Motor Learning in Speech Production. Neuroscience Letters, 591, 25–29. https://doi.org/10.1016/j.neulet.2015.02.012

Max, L., Wallace, M. E., & Vincent, I.Sensorimotor adaptation to auditory perturbations during speech: Acoustic and kinematic experiments. Paper presented at the International Congress of Phonetic Sciences, 1053-1056.

Mazzoni, P., & Krakauer, J. W. (2006). An Implicit Plan Overrides an Explicit Strategy during Visuomotor Adaptation. The Journal of Neuroscience: The Official Journal of the Society for Neuroscience, 26(14), 3642–3645. https://doi.org/10.1523/JNEUROSCI.5317-05.2006

McDougle, S. D., Bond, K. M., & Taylor, J. A. (2015). Explicit and Implicit Processes Constitute the Fast and Slow Processes of Sensorimotor Learning. The Journal of Neuroscience, 35(26), 9568–9579. https://doi.org/10.1523/JNEUROSCI.5061-14.2015

Neilson, M. D., & Neilson, P. D. (1987). Speech Motor Control and Stuttering: A Computational Model of Adaptive Sensory-Motor Processing. Speech Communication, 6, 325–333.

Nippold, M. A. (2002). Stuttering and Phonology: Is there an Interaction?. American Journal of Speech-Language Pathology, 11(2), 99–110. https://doi.org/10.1044/1058-0360(2002/011)

Oldfield, R. C. (1971). The Assessment and Analysis of Handedness: The Edinburgh Inventory. Neuropsychologia, 9(1), 97–113. https://doi.org/10.1016/0028-3932(71)90067-4

Packman, A., & Attanasio, J. S. (2004). Theoretical issues in stuttering. Psychology Press.

R Core Team. (2018). R: A Language and Environment for Statistical Computing [computer software]. Vienna, Austria:

Rabe, K., Livne, O., Gizewski, E. R., Aurich, V., Beck, A., Timmann, D., & Donchin, O. (2009). Adaptation to Visuomotor Rotation and Force Field Perturbation is Correlated to Different Brain Areas in Patients with Cerebellar Degeneration. Journal of Neurophysiology, 101(4), 1961–1971. https://doi.org/10.1152/jn.91069.2008

Raza, M. H., Amjad, R., Riazuddin, S., & Drayna, D. (2012). Studies in a Consanguineous Family Reveal a Novel Locus for Stuttering on Chromosome 16q. Human Genetics, 131(2), 311–313. https://doi.org/10.1007/s00439-011-1134-2

Riley, G. (1994). Stuttering Severity Instrument for Children and Adults. Doi,

Riley, G. (2009). SSI-4: Stuttering severity instrument

Robertson, E., & Miall, R. (1999). Visuomotor Adaptation during Inactivation of the Dentate Nucleus. Neuroreport, 10(5), 1029–1034. https://doi.org/10.1097/00001756-199904060-00025

Rossi, C., Chau, C. W., Leech, K. A., Statton, M. A., Gonzalez, A. J., & Bastian, A. J. (2019). The Capacity to Learn New Motor and Perceptual Calibrations Develops Concurrently in Childhood. Scientific Reports, 9(1), 9322. https://doi.org/10.1038/s41598-019-45074-6

Schlerf, J. E., Xu, J., Klemfuss, N. M., Griffiths, T. L., & Ivry, R. B. (2013). Individuals with Cerebellar Degeneration show Similar Adaptation Deficits with Large and Small Visuomotor Errors. Journal of Neurophysiology, 109(4), 1164–1173. https://doi.org/10.1152/jn.00654.2011

Schween, R., & Hegele, M. (2017). Feedback Delay Attenuates Implicit but Facilitates Explicit Adjustments to a Visuomotor Rotation. Neurobiology of Learning and Memory, 140, 124–133. https://doi.org/10.1016/j.nlm.2017.02.015

Semel, E. M., Wiig, E. H., & Secord, W. (2004). CELF 4. PsychCorp.

Sengupta, R., Shah, S., Gore, K., Loucks, T. M., & Nasir, S. M. (2016). Anomaly in Neural Phase Coherence Accompanies Reduced Sensorimotor Integration in Adults Who Stutter. Neuropsychologia, 93(Pt A), 242–250. https://doi.org/10.1016/j.neuropsychologia.2016.11.004

Shadmehr, R., Smith, M. A., & Krakauer, J. W. (2010). Error Correction, Sensory Prediction, and Adaptation in Motor Control. Annual Review of Neuroscience, 33(1), 89–108. https://doi.org/10.1146/annurev-neuro-060909-153135

Shahed, J., & Jankovic, J. (2001). Re-Emergence of Childhood Stuttering in Parkinson’s Disease: A Hypothesis. Movement Disorders: Official Journal of the Movement Disorder Society, 16(1), 114–118. https://doi.org/10.1002/1531-8257(200101)16:13.0.co;2-2

Shiller, D. M., Mitsuya, T., & Max, L. (2020). Exposure to Auditory Feedback Delay while Speaking Induces Perceptual Habituation but does Not Mitigate the Disruptive Effect of Delay on Speech Auditory-Motor Learning. Neuroscience, https://doi.org/10.1016/j.neuroscience.2020.07.041

Sitek, K. R., Cai, S., Beal, D. S., Perkell, J. S., Guenther, F. H., & Ghosh, S. S. (2016). Decreased Cerebellar-Orbitofrontal Connectivity Correlates with Stuttering Severity: Whole-Brain Functional and Structural Connectivity Associations with Persistent Developmental Stuttering. Frontiers in Human Neuroscience, 10, 190. https://doi.org/10.3389/fnhum.2016.00190

Smith, M. A., Ghazizadeh, A., & Shadmehr, R. (2006). Interacting Adaptive Processes with Different Timescales Underlie Short-Term Motor Learning. PLOS Biology, 4(6), e179. https://doi.org/10.1371/journal.pbio.0040179

Smith, M. A., & Shadmehr, R. (2005). Intact Ability to Learn Internal Models of Arm Dynamics in Huntington’s Disease but Not Cerebellar Degeneration. Journal of Neurophysiology, 93(5), 2809–2821. https://doi.org/10.1152/jn.00943.2004

Smits-Bandstra, S., & De Nil, L. (2009). Speech Skill Learning of Persons Who Stutter and Fluent Speakers Under Single and Dual Task Conditions. Clinical Linguistics & Phonetics, 23(1), 38–57. https://doi.org/10.1080/02699200802394914

Smits-Bandstra, S., & De Nil, L. F. (2007). Sequence Skill Learning in Persons Who Stutter: Implications for Cortico-Striato-Thalamo-Cortical Dysfunction. Journal of Fluency Disorders, 32(4), 251–278. https://doi.org/10.1016/j.jfludis.2007.06.001

Smits-Bandstra, S., De Nil, L. F., & Saint-Cyr, J. (2006a). Speech and Nonspeech Sequence Skill Learning in Adults Who Stutter. Journal of Fluency Disorders, 31(2), 116–136. https://doi.org/10.1016/j.jfludis.2006.04.003

Smits-Bandstra, S., De Nil, L., & Rochon, E. (2006b). The Transition to Increased Automaticity during Finger Sequence Learning in Adult Males Who Stutter. Journal of Fluency Disorders, 31(1), 22–40. https://doi.org/10.1016/j.jfludis.2005.11.004

Taylor, J. A., Krakauer, J. W., & Ivry, R. B. (2014). Explicit and Implicit Contributions to Learning in a Sensorimotor Adaptation Task. The Journal of Neuroscience: The Official Journal of the Society for Neuroscience, 34(8), 3023–3032. https://doi.org/10.1523/JNEUROSCI.3619-13.2014

Tong, C., & Flanagan, J. R. (2003). Task-Specific Internal Models for Kinematic Transformations. Journal of Neurophysiology, 90(2), 578–585. https://doi.org/10.1152/jn.01087.2002

Torchiano, M. (2018). Effsize - A Package for Efficient Effect Size Computation https://doi.org/10.5281/zenodo.1480624

Unicomb, R., Kefalianos, E., Reilly, S., Cook, F., & Morgan, A. (2020). Prevalence and Features of Comorbid Stuttering and Speech Sound Disorder at Age 4 Years. Journal of Communication Disorders, 84, 105976. https://doi.org/10.1016/j.jcomdis.2020.105976

Vallabha, G. K., & Tuller, B. (2002). Systematic Errors in the Formant Analysis of Steady-State Vowels. Speech Communication, 38(1), 141–160. https://doi.org/10.1016/S0167-6393(01)00049-8

Venkatakrishnan, A., Banquet, J. P., Burnod, Y., & Contreras-vidal, J. (2011). Parkinson’s Disease Differentially Affects Adaptation to Gradual as Compared to Sudden Visuomotor Distortions. Human Movement Science, 30(4), 760–769. https://doi.org/10.1016/j.humov.2010.08.020

Vorperian, H. K., & Kent, R. D. (2007). Vowel Acoustic Space Development in Children: A Synthesis of Acoustic and Anatomic Data. Journal of Speech, Language, and Hearing Research : JSLHR, 50(6), 1510–1545. https://doi.org/10.1044/1092-4388(2007/104)

Vorperian, H. K., Kent, R. D., Gentry, L. R., & Yandell, B. S. (1999). Magnetic Resonance Imaging Procedures to Study the Concurrent Anatomic Development of Vocal Tract Structures: Preliminary Results. International Journal of Pediatric Otorhinolaryngology, 49, 197–206. https://doi.org/10.1016/s0165-5876(99)00208-6

Vorperian, H. K., Kent, R. D., Lindstrom, M. J., Kalina, C. M., Gentry, L. R., & Yandell, B. S. (2005). Development of Vocal Tract Length during Early Childhood: A Magnetic Resonance Imaging Study. The Journal of the Acoustical Society of America, 117(1), 338–350. https://doi.org/10.1121/1.1835958

Vorperian, H. K., Wang, S., Chung, M. K., Schimek, E. M., Durtschi, R. B., Kent, R. D., Ziegert, A. J., & Gentry, L. R. (2009). Anatomic Development of the Oral and Pharyngeal Portions of the Vocal Tract: An Imaging Study. The Journal of the Acoustical Society of America, 125(3), 1666–1678. https://doi.org/10.1121/1.3075589

Webster, W. G. (1997). Principles of human brain organization related to lateralization of language and speech motor functions in normal speakers and stutterers. In W. Hustijn, H. F. M. Peters & P. van Lieshout (Eds.), Speech Production: Motor Control, Brain Research and Fluency Disorders. (pp. 119–139). Elsevier.

Webster, W. G., & Ryan, C. R. (1991). Task Complexity and Manual Reaction Times in People Who Stutter. Journal of Speech and Hearing Research, 34(4), 708–714. https://doi.org/10.1044/jshr.3404.708

Weiner, M. J., Hallett, M., & Funkenstein, H. H. (1983). Adaptation to Lateral Displacement of Vision in Patients with Lesions of the Central Nervous System. Neurology, 33(6), 766–772.

Werner, S., Bock, O., Gizewski, E. R., Schoch, B., & Timmann, D. (2010). Visuomotor Adaptive Improvement and Aftereffects are Impaired Differentially Following Cerebellar Lesions in SCA and PICA Territory. Experimental Brain Research.Experimentelle Hirnforschung.Experimentation Cerebrale, 201(3), 429–439. https://doi.org/10.1007/s00221-009-2052-6

Werner, S., Bock, O., & Timmann, D. (2009). The Effect of Cerebellar Cortical Degeneration on Adaptive Plasticity and Movement Control. Experimental Brain Research.Experimentelle Hirnforschung.Experimentation Cerebrale, 193(2), 189–196. https://doi.org/10.1007/s00221-008-1607-2

Williams, K. T. (2007). EVT-2: Expressive vocabulary test. Pearson Assessments.

Wolpert, D. M., & Miall, R. C. (1996). Forward Models for Physiological Motor Control. Neural Networks : The Official Journal of the International Neural Network Society, 9(8), 1265–1279.

Wolpert, D. M., & Flanagan, J. R. (2001). Motor Prediction. Current Biology: CB, 11(18), 729. https://doi.org/10.1016/s0960-9822(01)00432-8

Wolpert, D. M., & Flanagan, J. R. (2009). Forward models. The Oxford Companion to Consciousness (pp. 295-296). OUP Oxford.

Wu, J. C., Maguire, G., Riley, G., Fallon, J., LaCasse, L., Chin, S., Klein, E., Tang, C., Cadwell, S., & Lottenberg, S. (1995). A Positron Emission Tomography [18F]Deoxyglucose Study of Developmental Stuttering. Neuroreport, 6(3), 501–505.

Wu, J. C., Maguire, G., Riley, G., Lee, A., Keator, D., Tang, C., Fallon, J., & Najafi, A. (1997). Increased Dopamine Activity Associated with Stuttering. Neuroreport, 8(3), 767–770. https://doi.org/10.1097/00001756-199702100-00037

Wymbs, N. F., Ingham, R. J., Ingham, J. C., Paolini, K. E., & Grafton, S. T. (2013). Individual Differences in Neural Regions Functionally Related to Real and Imagined Stuttering. Brain and Language, 124(2), 153–164. https://doi.org/10.1016/j.bandl.2012.11.013

Zelaznik, H. N., Smith, A., Franz, E. A., & Ho, M. (1997). Differences in Bimanual Coordination Associated with Stuttering. Acta Psychologica, 96(3), 229–243. https://doi.org/10.1016/s0001-6918(97)00014-0

